# Design of intrinsically disordered protein variants with diverse structural properties

**DOI:** 10.1101/2023.10.22.563461

**Authors:** Francesco Pesce, Anne Bremer, Giulio Tesei, Jesse B. Hopkins, Christy R. Grace, Tanja Mittag, Kresten Lindorff-Larsen

**Affiliations:** Structural Biology and NMR Laboratory, The Linderstrøm-Lang Centre for Protein Science, Department of Biology, University of Copenhagen, Copenhagen, Denmark; Department of Structural Biology, St. Jude Children’s Research Hospital, Memphis, TN 38105, USA; BioCAT, Department of Physics, Illinois Institute of Technology, Chicago, IL, USA

## Abstract

Intrinsically disordered proteins (IDPs) perform a wide range of functions in biology, suggesting that the ability to design IDPs could help expand the repertoire of proteins with novel functions. Designing IDPs with specific structural or functional properties has, however, been difficult, in part because determining accurate conformational ensembles of IDPs generally requires a combination of computational modelling and experiments. Motivated by recent advancements in efficient physics-based models for simulations of IDPs, we have developed a general algorithm for designing IDPs with specific structural properties. We demonstrate the power of the algorithm by generating variants of naturally occurring IDPs with different levels of compaction and that vary more than 100 fold in their propensity to undergo phase separation, even while keeping a fixed amino acid composition. We experimentally tested designs of variants of the low-complexity domain of hnRNPA1 and find high accuracy in our computational predictions, both in terms of single-chain compaction and propensity to undergo phase separation. We analyze the sequence features that determine changes in compaction and propensity to phase separate and find an overall good agreement with previous findings for naturally occurring sequences. Our general, physics-based method enables the design of disordered sequences with specified conformational properties. Our algorithm thus expands the toolbox for protein design to include also the most flexible proteins and will enable the design of proteins whose functions exploit the many properties afforded by protein disorder.

## Introduction

Intrinsically disordered proteins and regions (from here collectively termed IDPs) (***Uversky and Dunker, 2010***) represent a diverse class of proteins that carry out a wide range of functions (Van Der Lee et al., 2014) and are characterized by extreme but often non-random structural heterogeneity. Their distinct amino acid composition and sequences (***Uversky et al., 2000***) differ from those of natively folded proteins, and prevent the formation of stable folded conformations. Thus, IDPs are best described by ensembles of heterogeneous conformations that interconvert rapidly (***Mittag and Forman-Kay, 2007***; ***Thomasen and Lindorff-Larsen, 2022***). The disordered and dynamic nature of IDPs is often central for their biological and biochemical functions. They can be linkers separating functional domains, regulating the interaction between the latter (***Li et al., 2018***), or they can play roles as spacers that impair undesirable protein-protein interactions (***Santner et al., 2012***; ***Jamecna et al., 2019***). IDPs are often involved in mediating molecular interactions including via so-called short-linear motifs (***Davey et al., 2012***), and their large capture radius may give rise to faster binding kinetics compared to that of folded proteins (***Shoemaker et al., 2000***). Thus, IDPs are for example commonly found in signaling molecules (***Wright and Dyson, 2015***) and transcription factors (***Liu et al., 2006***). Furthermore, the interactions within and between IDPs and other biomolecules have emerged as an important factor in the spatial organization of cellular matter. Through their ability to form multivalent interactions, IDPs can aid in or drive the formation of membraneless organelles, which typically consist of a wide range of biomolecules and compartmentalize many biological processes (***Banani et al., 2017***; ***Mittag and Pappu, 2022***). In vitro, many IDPs have been shown to undergo a phase separation (PS) process that leads to the co-existence of a protein-rich dense phase that separates from a dilute phase when the concentration of the protein reaches the so-called saturation concentration (*c*_sat_) (***Mittag and Pappu, 2022***). Thus, at concentrations above *c*_sat_, the protein may be found both in a dilute phase, and a co-existing dense phase that macroscopically may appear liquid-like and at the molecular level may behave as a viscoelastic fluid (***Mittag and Pappu, 2022***; ***Alshareedah et al., 2023***).

Similarly to the long-lasting quest for predicting the native structure of folded proteins from their sequences (***Kuhlman and Bradley, 2019***), a field which has recently witnessed substantial advances (***Jumper et al., 2021***; ***Baek et al., 2021***; ***Lin et al., 2023***), there is interest in understanding the sequence determinants for the conformational properties of IDPs (***Uversky et al., 2000***; ***Marsh and Forman-Kay, 2010***; ***Das et al., 2015***; ***Cohan et al., 2019***) and how these are related to their function (***Zarin et al., 2021***; ***Tesei et al., 2023***). For both folded and disordered proteins, the ability to predict structure(s) from sequences may help infer its functional properties. Accurate structure prediction may also support or sometimes replace the need for experimental studies of protein structure. Finally, rapid structure prediction enables proteome-wide analyses and can aid in protein design.

In parallel with our continuously improving ability to predict structures of folded proteins, there has been substantial development in our ability to design sequences that fold into specific three-dimensional folded structures (***Pan and Kortemme, 2021***; ***Woolfson, 2021***; ***Goverde et al., 2023***). Given the multitude of functions and properties of IDPs, there would be a great potential in designing IDPs with targeted properties. Such proteins could potentially find applications in designing linkers in multi-domain enzymes (***Van Rosmalen et al., 2017***), signalling molecules, or using IDPs as biomaterials (***Dzuricky et al., 2018***). In contrast to the developments for folded proteins, our ability to design IDPs with specific properties remains more limited. This is because characterizing and predicting the structural properties of IDPs is a complicated task, and because we know less about the sequence-ensemble relationships for IDPs. The native structure of folded proteins can be experimentally determined at atomic resolution, and the availability of many high-resolution structures has been one key driving force to understand and predict how sequences encode structures (***Jumper et al., 2021***). On the other hand, characterizing the ensemble of conformations that an IDP adopts generally requires the integration of experiments and simulation methods (***Mittag and Forman-Kay, 2007***; ***Thomasen and Lindorff-Larsen, 2022***). Collecting and interpreting such data is, however, difficult and often ambiguous, and as a consequence there are only limited examples of detailed structural characterizations (Lazar et al., 2021). Thus, there are still many open questions about how the sequence of an IDP translates into a structural ensemble and function (Lindorff-Larsen and Kragelund, 2021). Despite these limitations, a number of rules have emerged that govern the local and global conformational properties of IDPs. For example, the content (Müller-Späth et al., 2010) and patterning (***Das and Pappu, 2013***) of charged residues has been related to the global expansion of an IDP in solution (***Tesei et al., 2023***; ***Lotthammer et al., 2023***), as well as their propensity to undergo PS (***Lin and Chan, 2017***; ***Schuster et al., 2020***; ***Bremer et al., 2022***). Similarly, hydrophobicity, and in particular the number and patterning of aromatic residues, influences the compaction of an IDP and its propensity to phase separate (***Zheng et al., 2020***; ***Martin et al., 2020***; ***Holehouse et al., 2021***).

A number of different approaches have recently enabled the development of accurate, yet highly computationally-efficient models for molecular simulations of the global conformational properties of IDPs (***Shea et al., 2021***; ***Tesei et al., 2021***; ***Dannenhoffer-Lafage and Best, 2021***; ***Regy et al., 2021***; ***Joseph et al., 2021***; ***Tesei and Lindorff-Larsen, 2022***). These simulation methods make it possible to use a physics-based coarse-grained model to predict conformational ensembles from sequences on time-scales that are compatible with screening large number of sequences, e.g. all IDPs in the human genome (***Tesei et al., 2023***). Building on these developments, we here present an algorithm to generate sequences of IDPs with pre-defined conformational properties. The cen- tral idea is to search sequence space and to use efficient coarse-grained simulations to link each sequence to conformational properties. Specifically, we use the CALVADOS model, that has been optimized by targeting small-angle X-ray scattering (SAXS) and paramagnetic relaxation enhance- ment NMR experiments on IDPs in solution (***Tesei et al., 2021***), and which has been extensively validated using independent experimental data (***Tesei et al., 2023***). In some aspects our work builds on previous work using genetic algorithms (***Zeng et al., 2021***; ***Lichtinger et al., 2021***), but we show how our design method enables large-scale exploration of the sequence-structure space and validate the results experimentally.

We begin by studying four IDPs with different sequence compositions and characteristics. Start- ing from each sequence, we design new sequences with different levels of compaction while keep- ing the amino acid composition constant. The results show that—even with the restriction of hav- ing a fixed amino acid composition—it is possible to achieve conformational ensembles with highly diverse properties. We show that this is mainly, but not solely, due to differences in the patterning of charges. We used the low complexity domain of hnRNPA1 (hereafter A1-LCD), to study the rela- tionship between sequence patterning, single-chain properties, and the propensity to undergo PS. We selected five variants of A1-LCD for experimental characterization, and find good agreement between the experiments and predictions. Together, our results show that the algorithm that we have developed is efficient and can be used to design IDP sequences with novel properties. The algorithm is fully general, and can therefore also be used to design sequences with varying amino acid composition and for other target properties than chain expansion.

## Results

### Algorithm to design novel IDPs

To design IDP sequences with specific conformational properties, it is necessary to be able to pre- dict these properties from sequences accurately and rapidly. Therefore, the first question that we address is whether it is possible to use state-of-the-art simulation-based approaches to develop a generalizable method for IDP design. Very recent work has established efficient machine-learning- based methods to predict average conformational properties from sequences (***Tesei et al., 2023***; ***Lotthammer et al., 2023***), but these methods do not predict full conformational ensembles and have not been tested experimentally on novel sequences. Instead, we used a simulation-based approach where we employ a coarse-grained model to generate a conformational ensemble for a given sequence (Fig. 1).

**Figure 1.**
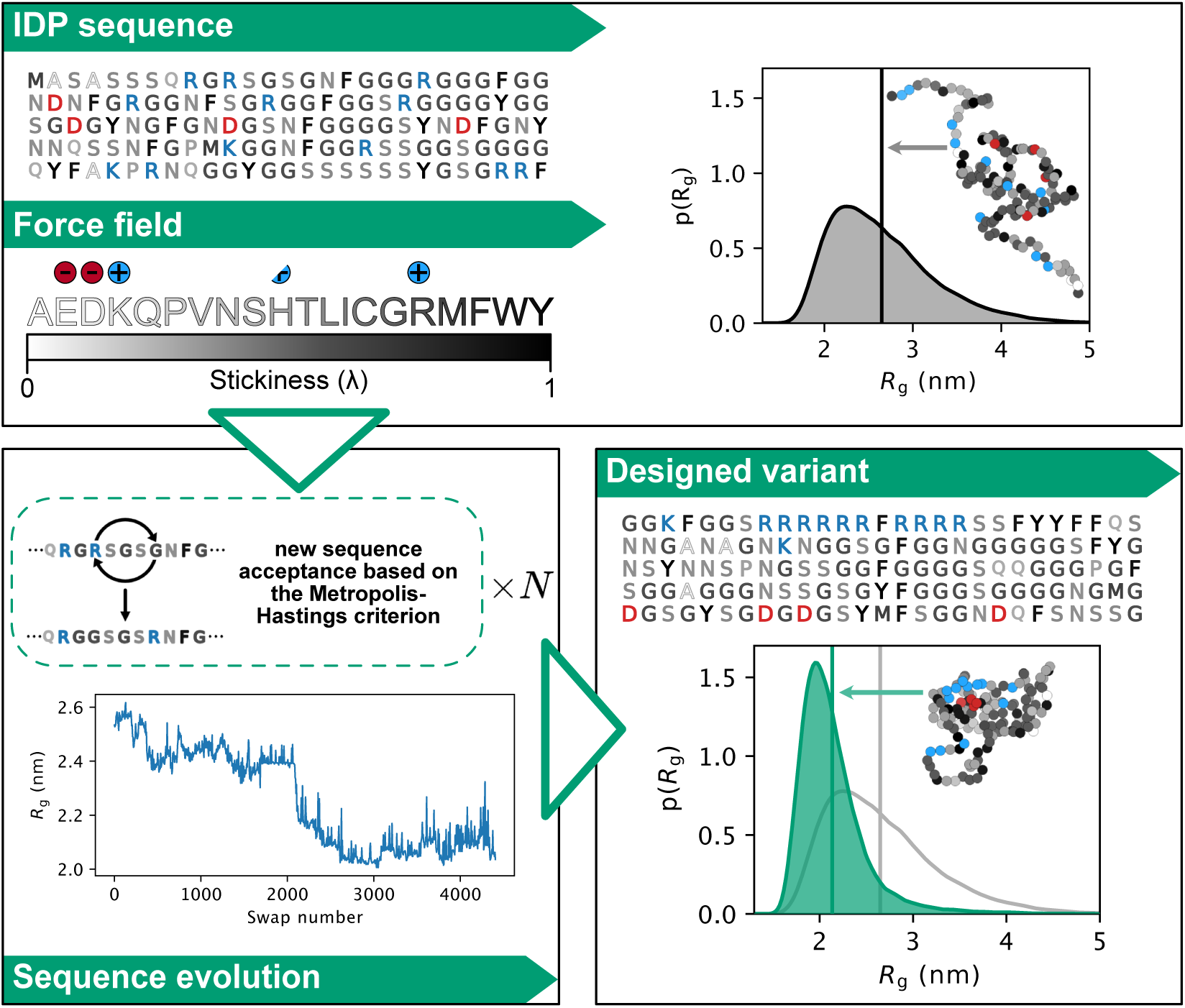
Outline of our algorithm for designing sequences of IDPs with targeted conformational properties. As starting point, we here use naturally occurring IDP sequences, though this is not a requirement of the approach. We use MD simulations with the coarse-grained CALVADOS force field to describe the IDPs and to generate a conformational ensemble. New sequences are proposed through a Markov chain Monte Carlo scheme. We evolve the sequences by consecutive swaps in positions between two randomly selected residues, and evaluate whether the sequences get closer or further away from the design target—here chain compaction. During sequence optimization, we calculate the conformational properties for a given sequence either by direct simulations or through alchemical calculations that rely on conformational ensembles of previously sampled sequences. The conformations shown have the same radius of gyration as the average of the conformational ensemble.

We combine coarse-grained molecular dynamics (MD) simulations using the CALVADOS model (***Tesei et al., 2021***) with alchemical free-energy calculations in an algorithm that sequentially gener- ates new sequences and characterizes their conformational ensembles in a time-efficient manner. While MD simulations with a coarse-grained model can rapidly produce conformational ensem- bles from which structural features can be directly calculated, screening a large number of different IDPs sequentially with only MD simulations would still be computationally difficult. Alchemical free- energy calculations, on the other hand, can predict conformational properties of newly proposed sequences from conformational ensembles generated by simulations of different sequences. Our algorithm thus combines simulations and alchemical free-energy calculations in an optimization process that in some ways is analogous to what has been proposed in the context of force field optimization (***Norgaard et al., 2008***; ***Orioli et al., 2020***; Köfinger and Hummer, 2021).

While the overall sequence composition of an IDP is known to affect its conformational proper- ties (***Tesei et al., 2023***), we here aimed at exploring the more subtle and difficult-to-extract effects of sequence patterning (***Das and Pappu, 2013***; ***Das et al., 2015***; ***Sherry et al., 2017***; ***Beveridge et al., 2019***; ***Martin et al., 2020***; ***Cohan et al., 2021***). Therefore, we apply our design algorithm to gener- ate sequences of IDPs with diverse structural properties while preserving the overall amino acid composition. In this way we also test and possibly expand our understanding of how the pattern- ing of specific residues in a sequence influences its conformational properties. Early pioneering work focused on the role of charge patterning on conformational properties and propensity to phase separate (***Das and Pappu, 2013***; ***Das et al., 2016***; ***Lin and Chan, 2017***; ***Schuster et al., 2020***). Other studies have linked the number and patterning of amino acids, in particular aromatic and arginine residues, to both conformational and phase properties (***Wang et al., 2018***; ***Martin et al., 2020***; ***Holehouse et al., 2021***; ***Bremer et al., 2022***).

Nonetheless, even restricting the sequence space to sequences of fixed composition, the num- ber of possible sequences is enormous; for example, there are ca. 1.8×10^127^ unique sequences with the amino acid composition of the disordered domain of the fused in sarcoma (FUS) protein. Thus, sampling even a tiny part of this space is unfeasible. To circumvent this problem, our algorithm drives the exploration of the sequence space towards sequences resulting in the target conforma- tional property. This is achieved via a Markov chain Monte Carlo (MCMC) sampling scheme that iteratively generates sequence variants and predicts their conformational properties (through MD simulations and alchemical free-energy calculations) in search of specific arrangements of amino acids that determine a certain structural feature (see Methods for a more detailed description of the algorithm and its components).

To exemplify and demonstrate the power of our algorithm we generate variants of IDPs with either increased or decreased chain expansion, measured by their radius of gyration (*R*_g_), while keeping a fixed amino acid composition. To this aim, at each iteration the algorithm swaps the positions of two randomly selected residues to generate a variant (from hereon called a swap variant). We compare the *R*_g_ before and after the swap (evaluated either from MD simulations or alchemical free-energy calculations), and the Monte Carlo move is accepted or rejected based on the Metropolis-Hastings criterion (Fig. 1). Although we here have focused on the difficult problem of changing conformational properties while keeping a fixed amino acid composition, the algorithm is versatile and other criteria can be used to propose changes in the sequences (e.g. single point mutations without keeping a fixed amino acid composition) as well as selecting for other structural features than the *R*_g_.

### Design of IDPs with conformational ensembles that vary in compaction

The second question that we address is: Starting from a natural IDP, how much more compact or expanded can it become when only changing the positions of the amino acids in its sequence? To answer this question, we selected four IDPs with different sequence compositions: *α*-Synuclein (*a*Syn), and the low complexity domain from hnRNPA1 (A1-LCD), the prion-like domain in FUS (FUS- PLD) and the R-/G-rich domain of the P granule protein LAF-1 (LAF-1-RGG) (Fig. 2a). We used our sequence design algorithm in a simulated annealing protocol to let the sequences evolve in search of amino acid arrangements that result in more compact ensembles. The results show that we can generate sequence permutations of *α*Syn, A1-LCD and LAF-1-RGG, that are substantially more compact than the wild-type sequence (Fig. 2b, green lines). In contrast, for FUS-PLD we only find variants that are modestly more compact than the wild-type protein. To demonstrate that the algorithm can also find sequences of increased expansion, we began from the compact designs and instead targeted greater *R*_g_ values. For *α*Syn, A1-LCD and LAF-1-RGG we find that the algorithm quickly generates sequences with wild-type-like dimensions (Fig. 2b, orange lines). Interestingly, in all cases the algorithm only finds sequences that are modestly more expanded than the wild-type sequence although the algorithm was tuned to expand the protein as much as possible. We repeated these calculations starting also from the wild-type sequences and reached similar results (Fig. S1).

**Figure 2.**
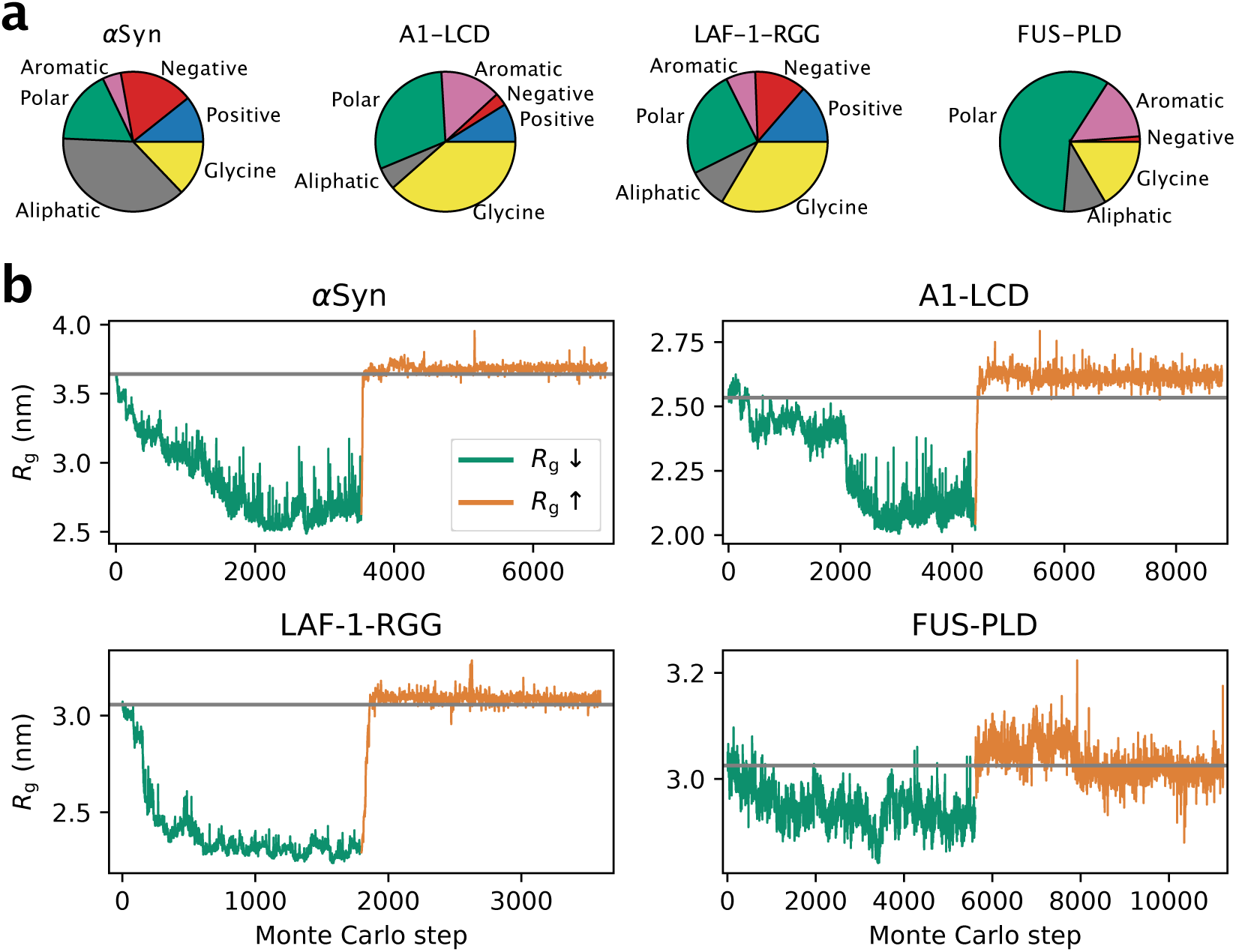
(a) Pie chart of the sequence composition of *α*Syn, A1-LCD, LAF-1-RGG and FUS-PLD. Amino acids are grouped as negative (D, E), positive (R, K), aromatic (Y, W, F), polar (S, T, N, Q, H, C), aliphatic (A, V, I, L, M, P) and glycine. (b) Design of compact (green lines) and expanded (orange lines) variants for *α*Syn, A1-LCD, LAF-1-RGG and FUS-PLD. Each accepted Monte Carlo step thus gives rise to a sequence that differs from the previous by the position of the two swapped residues. Each Monte Carlo step therefore corresponds to a different sequence, whose ensemble averaged *R*_g_ is evaluated by either MD simulations or alchemical free-energy calculations. The grey horizontal line indicates the *R*_g_ of the wild-type sequence.

### Sequence features that determine the compaction of the designs

In the calculations above, we observed that while thousands of swap moves are required for the algorithm to reach the most compact ensembles, a much smaller number of moves was required to recover sequences with wild-type-like dimensions (Fig. 2b). As the moves swap two randomly selected positions, we speculate that there is an entropic barrier in sequence space in finding the arrangement of amino acids that determines compact ensembles. This suggests compaction is driven by some kind of specific ordering of the amino acid sequences. The next question we ad- dressed was therefore: What are the sequence determinants of IDP compaction in the generated sequences? As described above, we were able to generate substantially more compact variants for *a*Syn, A1-LCD and LAF-1-RGG, but not for FUS-PLD. We therefore aimed to identify which se- quence features led to this compaction, and assessed if the same features were responsible in all three cases. We calculated a number of sequence features for the variants of *α*Syn, A1-LCD and LAF-1-RGG and examined the correlation with the *R*_g_ (Figs. 3a and S2). In all cases, we observe a strong correlation between the patterning of the charged amino acid residues, as captured by the *K* parameter (***Das and Pappu, 2013***) (Fig. 3a), and chain dimensions. The *K* parameter captures whether the positively and negatively charged residues are well mixed together (low *κ*) or whether they tend to be found in blocks of like charges (high *κ*) (***Das and Pappu, 2013***). For all three proteins we observe that the positively charged residues tend to be clustered in the N-terminal third of the sequence and the negatively charged residues in the C-terminal third as the sequences get increasingly compact during the sequence design (Fig. 3b). Since the N-terminus carries a positive charge, and the C-terminus carries a negative charge, it is likely that the termini contribute to the overall charge segregation. We stress that we did not directly drive this charge clustering during the sequence design algorithm, but that the analysis shows that clustering of the charges occurs as the algorithm explores sequence space to generate compact structures. The formation of charge-clustered sequences is in line with the hypothesis above of an ‘entropic bottleneck’ during the sequence design, and that it is easier to disrupt such patterns than to generate them by randomly swapping amino acid residues.

**Figure 3.**
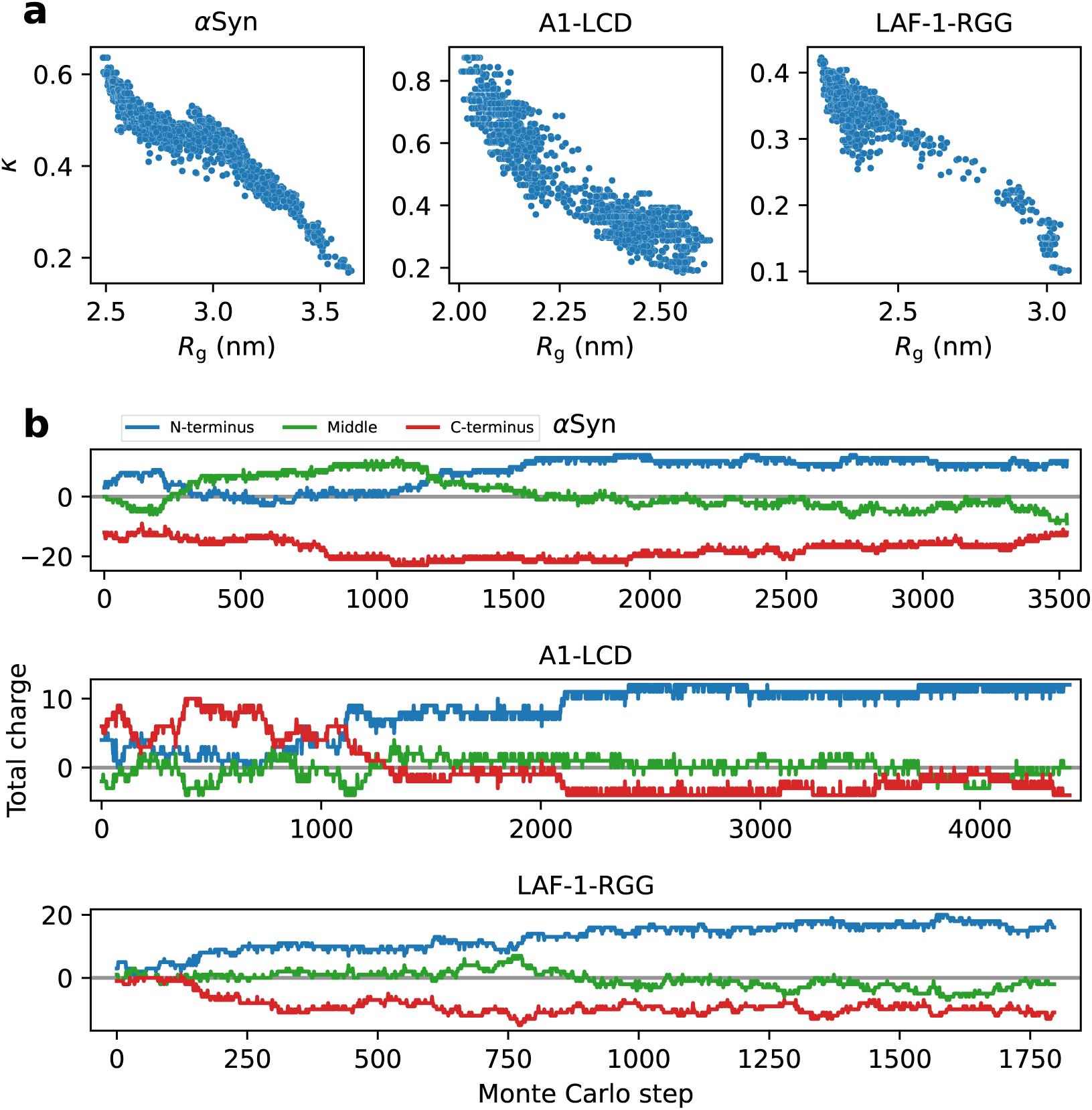
(a) Correlation between *R*_g_ and *K* (a high *K* indicates segregated clusters of residues with the same charge, a low *K* indicates that charges are well mixed along the sequence). (b) We divided the sequences of *a*Syn, A1-LCD and LAF-1-RGG into three sections covering the N-terminal third (blue), the middle third (green), and the C-terminal third (red) of the sequence, and calculated the total charge in each of these sections.

We also examined other sequence features including patterning of aromatic and hydrophobic residues, and found that they generally have a weaker correlation with the *R*_g_ (Fig. S2). For LAF- 1-RGG we, however, found that the patterning of hydrophobic residues may also contribute to compaction similarly to the patterning of charges (Fig. S2). This suggests that while charge patterning captures most of the variation in compaction of the permuted sequences, it is difficult to find individual sequence descriptors that fully explain the chain dimensions of IDPs, and that combinations of features may be needed to predict compaction (***Cohan et al., 2021***; ***Tesei et al., 2023***; ***Lotthammer et al., 2023***; ***Chao et al., 2023***). The importance of charge patterning also helps to explain why we were not able to obtain swap variants of FUS-PLD that are more compact than the wild-type, since FUS-PLD has only two negatively charged and no positively charged residues (Fig. 2a).

### Relating sequence, compaction and propensity to phase separate for the designs

Theory, simulations and experiments show that the compaction of an IDP is related to its propensity to self-associate and to undergo different forms of phase transitions (***Choi et al., 2020***). Conceptually, this can be understood by the fact that the intramolecular interactions that drive sequence compaction are the same as the intermolecular interactions that drive self-association and phase separation. It would be useful to be able to design proteins with predefined propensities to undergo phase separation and participate in the formation of biomolecular condensates. Building on previous work in this area (***Zeng et al., 2021***; ***Lichtinger et al., 2021***), the fourth question that we sought to answer is: Are the changes in single-chain compaction of the designed swap variants accompanied by a change in their propensity to phase separate? To examine this question we chose to study A1-LCD in more detail because the relationship between sequence and phase separation of A1-LCD has been studied extensively by experiments, theory and simulations (***Martin et al., 2020***; ***Tesei et al., 2021***; ***Bremer et al., 2022***; ***Maristany et al., 2023***).

To improve statistics, we performed nine additional runs of the design algorithm to generate a larger and more diversified pool of A1-LCD variants with different levels of compaction (Fig. S3). We then grouped these sequences by their *R*_g_ (in bins of 0.05-nm width), clustered the sequences (see Supplementary material), and use the centroid of each cluster for further analyses. In this way we remove sequences that are very similar to each other (there are many similar sequences within each run of sequence design since the design algorithm evolves sequences by consecutive position swaps of two residues) and only use one representative sequence for each cluster. We then performed 1- *µ*s simulations of each centroid sequence to re-evaluate their *R*_g_. We do this to validate the accuracy of the alchemical free-energy calculations in predicting the *R*_g_ of variants proposed by the design algorithm. In line with preliminary tests (Fig. S4, see Methods), we find an average error on the predicted *R*_g_ values of 1.5% (Fig. S5). We then re-binned the centroids based on the *R*_g_ from simulations, and for each bin we selected up to 15 sequences that are diverse in the patterning of charged and aromatic residues. In this way, we selected 120 A1-LCD swap variants (including the wild type) with diverse sequence features and compaction (Fig. 4a,b). Of the 119 swap variants, 113 have less than 30% sequence identity to the wild-type protein (Fig. S6).

**Figure 4.**
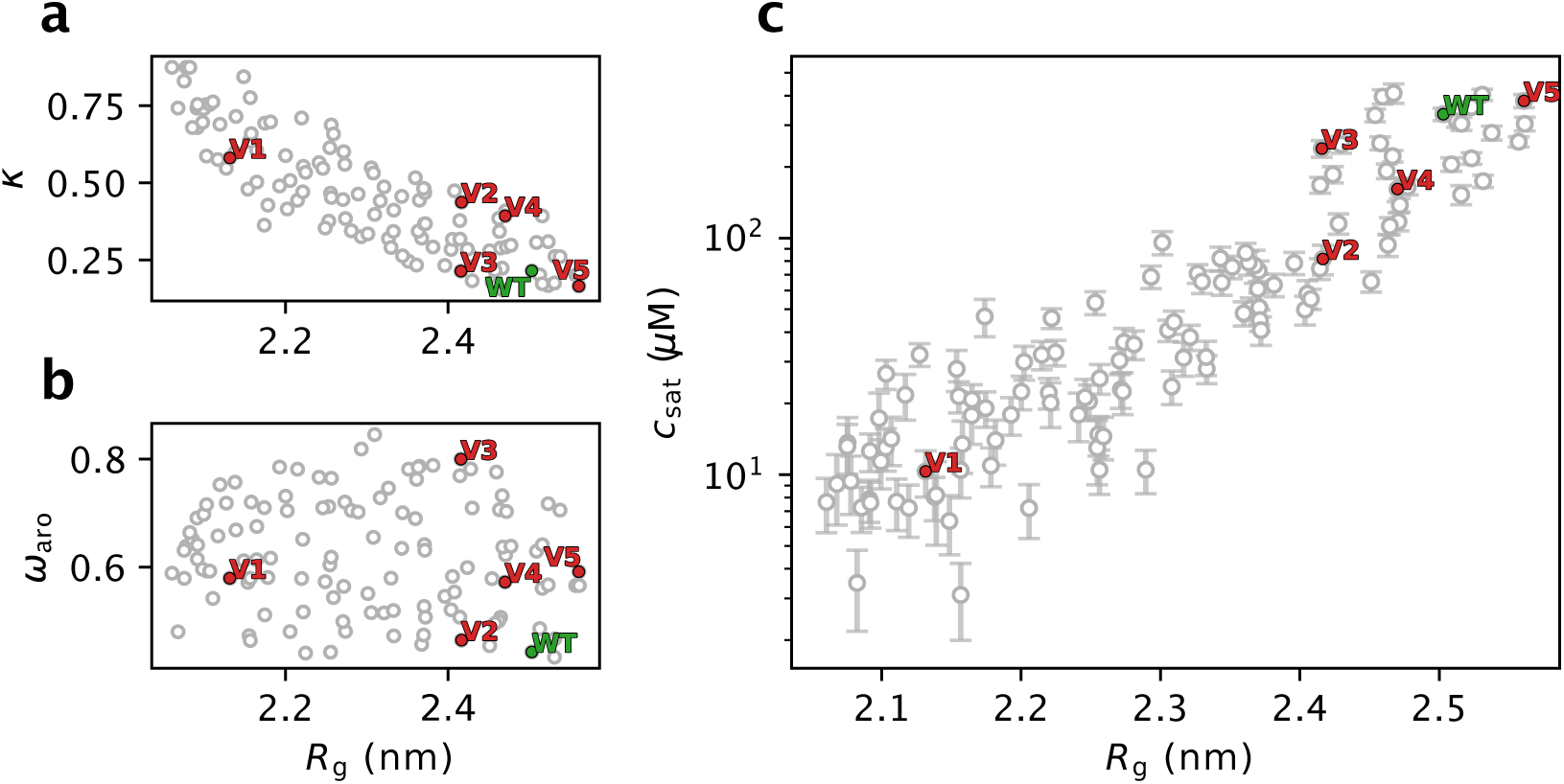
Characterization of the 119 A1-LCD swap variants selected by designing for more compact conformational ensembles and the wild-type (WT) A1-LCD. We show the relationship between *R*_g_ and (a) *κ*, (b) *cv*_aro_ (patterning of aromatic residues; a high *cv*_aro_ indicates clustering of aromatic residues), (c) the *c*_sat_ calculated from simulations of 100 chains in slab geometry. We highlight the WT sequence of A1-LCD in green and five variants selected for experimental characterization in red. Error bars of the average *R*_g_ are not shown as their size is negligible.

To examine the propensity of the designed A1-LCD variants to phase separate, we ran simulations of these variants (one at a time) consisting of 100 copies in a ‘slab’ geometry and estimate their *c*_sat_ from the concentration of the dilute phase in the simulation box (***Dignon et al., 2018***). As previously observed for a model system (***Lin and Chan, 2017***), we find a logarithmic relationship between *R*_g_ and *c*_sat_, with compact variants showing a stronger propensity to PS (low *c*_sat_), and expanded variants showing a weaker propensity for PS (high *c*_sat_) (Fig. 4c). Despite this expected correlation between single-chain properties and the propensity to phase separate, we find some sequences with similar *R*_g_ values whose *c*_sat_ values differ by up to one order of magnitude. This observation suggests that while the single chain behaviour can be very similar, other features encoded in the sequences can cause diversity in the PS properties. Overall, this correlation between *R*_g_ and *c*_sat_ further supports a strong link between single-chain properties and PS propensity that can be used to extrapolate PS propensity from single chain compaction, but also suggests that other sequence features that do not substantially change the single-chain *R*_g_ might have a role in PS.

### Experimental characterization of A1-LCD variants

Above we have described an approach to design IDPs and examine how the arrangement of amino acids in the primary sequences can influence their behaviour. While the coarse-grained model that we use in our algorithm (***Tesei et al., 2021***) has been extensively validated on naturally occurring proteins and variants thereof (***Tesei et al., 2023***), it has not been used as a generative model and tested on novel, designed sequences. We thus asked whether the accuracy of CALVADOS for pre- dicting *R*_g_ and *c*_sat_ for natural proteins also extends to sequences that show little sequence identity to natural proteins and, for example, show substantial charge patterning. Thus, a fifth question that we asked was: How accurate are our computational predictions of chain compaction and propensity to phase separate for the designed variants?

We therefore sought to test our predictions by experiments. We focused our experiments on fifteen swap variants of A1-LCD, selected from the 120 sequences analysed above, that represent a range of compaction and sequence properties. We focused on A1-LCD since the wild-type protein is already relatively compact and because its propensity to phase separate is rather strong for a protein of its length (***Martin et al., 2020***; ***Bremer et al., 2022***). Thus, we speculated that the ability to make it even more compact and endow it with lower *c*_sat_ without changing the amino acid composition would be a powerful test of our design algorithm and the CALVADOS model.

Out of the fifteen variants that we selected, we successfully expressed and purified five variants (red points in Fig. 4 and S7) and the wild-type A1-LCD protein. We ran new simulations of the se- lected variants under the conditions of the experiments and including a glycine-serine pair at the N-terminus that is present in the experimental constructs (Table S1). We name these variants V1 to V5, sorted by their calculated *R*_g_, with V1 predicted to be the most compact and most strongly phase separating variant, with a strong segregation of positive and negative charges at the termini (Fig. 5a). We induced phase separation by adding 150 mM NaCl and visualized the resulting con- densates by differential interference contrast (DIC) microscopy. We observed that all variants form condensates, and show some diversity in their morphology (Fig. 5b). We measured the *c*_sat_ of the five variants and the wild-type and compared the experimental results with those predicted from multi-chain simulations. We find a high correlation between predicted and observed values of *c*_sat_ (Fig. 5c), with the only outlier being V5, which is the sole variant expected to be more expanded than the WT (Fig. 5b). To investigate possible reasons for the discrepancy in PS propensity of V5 we ran additional simulations. The calculated *c_sat_* values that we compare to experiments (Fig. 5c) are averages over the *c_sat_* values calculated from three independent simulations. We obtained compa- rable results from the three independent replicates, demonstrating that the differences are not due to lack of convergence of the simulations (Fig. S8). We also ran simulations with different se- tups: one with twice as many chains to address potential finite size effects, and another with the updated CALVADOS 2 model (***Tesei and Lindorff-Larsen, 2022***). All three simulation setups gave comparable values for *c*_sat_ (Fig. S8).

**Figure 5.**
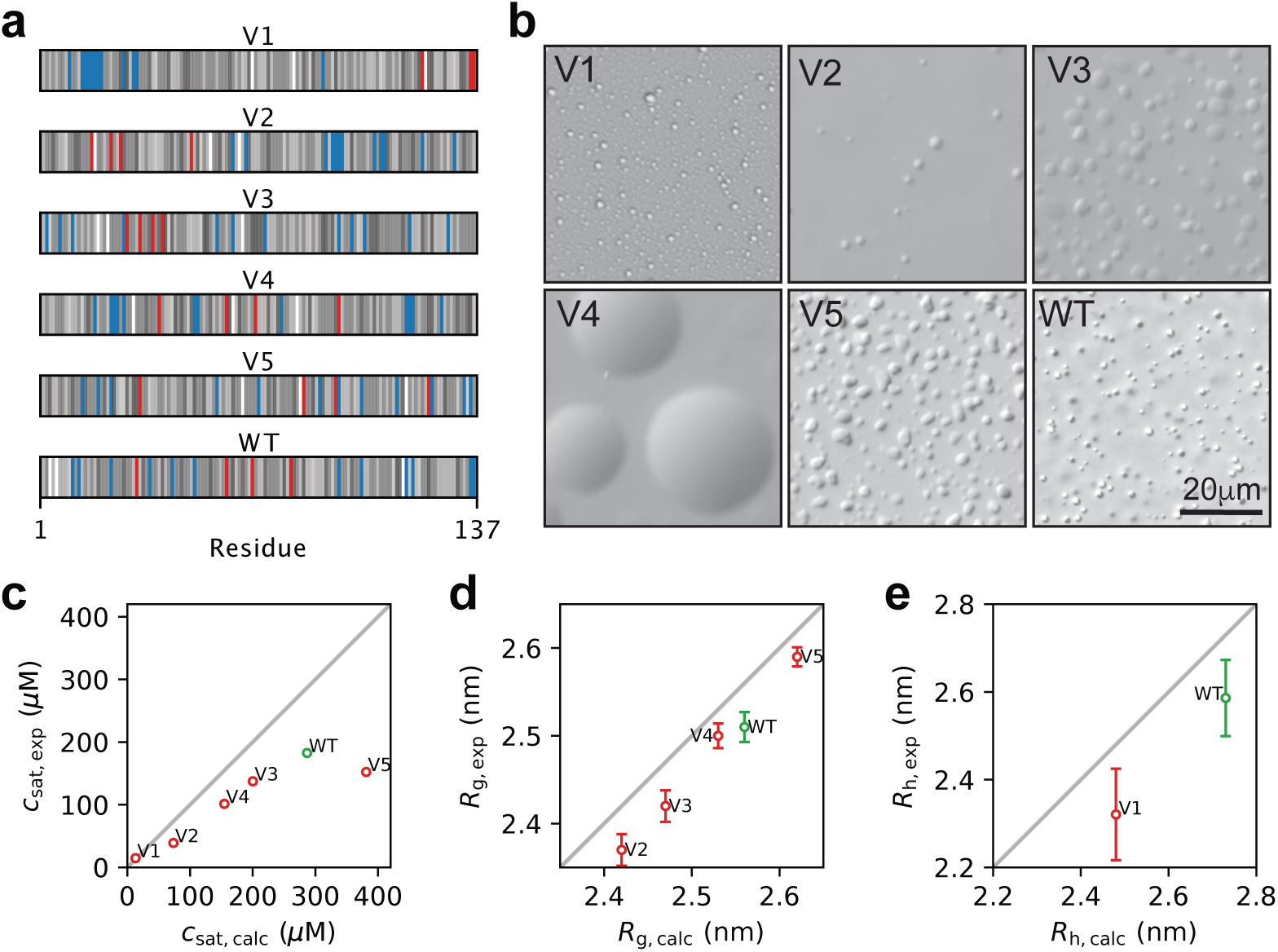
Experimental characterization of wild-type A1-LCD and five designed variants. (a) Diagram of the arrangement of amino acids in A1-LCD and the five design variants. Negative and positive charges are coloured respectively in red and blue. The neutral residues are coloured by a grey scale that reflects their hydrophobicity (corresponding to the *-1* parameter in CALVADOS), with the least hydrophobic residues in white and the most hydrophobic residues in black. (b) Visualization of condensates of wild-type A1-LCD and the five variants by DIC microscopy; these images are only meant to illustrate the formation of condensates and not necessarily differences between the variants. (c) Comparison of experimental and calculated values of *c*_sat_ at 298 K. (d) Comparison of experimental and calculated values of *R*_g_ for wild-type A1-LCD and V2–V5. (e) Comparison of experimental and calculated values of *R*_h_ at 304 K for wild-type A1-LCD and V1. Error bars whose sizes are comparable to that of the markers are not shown.

We used previously described methods to measure SAXS data for proteins close to the solubility limit (***Martin et al., 2021***) to test our predictions of sequence compaction. Like for *c*_sat_, we find a high correlation between the *R*_g_ values derived from SAXS and those from simulations (Fig. 5d), and a good agreement between the experimental and calculated SAXS curves with *x*^2^ values around 1–2 (Fig. S9). Given the low *c*_sat_ of V1 (15 *µ*M), we were not able to obtain a sufficient signal-to-noise ratio at a protein concentration below *c*_sat_. We instead turned to diffusion NMR experiments at low protein concentrations to measure the hydrodynamic radius (*R*_h_) of V1 and wild-type A1-LCD. We thus acquired NMR data for wild type A1-LCD and V1 at 307 K, where the measured *c*_sat_ of V1 is 34 *µ*M (compared to 15 *µ*M at 298 K). At this temperature, we find that V1 is substantially more compact than wild-type A1-LCD (Fig. 5e). We note that for both *R*_g_ and *R*_h_ there appears to be a small, but systematic, offset between the predicted and experimentally determined values. Some of these differences may indicate remaining errors in the CALVADOS force field, but may also reflect uncertainty in how *R*_g_ and *R*_h_ are estimated from experiments and simulations (***Henriques et al., 2018***; ***Pesce and Lindorff-Larsen, 2021***; ***Pesce et al., 2022***; ***Tranchant et al., 2023***), and we also note the high agreement between calculated and experimental SAXS data (Fig. S9).

We find that both simulations and experiments show that V3 is more compact than V4 (Fig. 5d), while V4 has a lower *c*_sat_ than V3 (Fig. 5c). Previously it has been shown that changes in the formal net charge may break the correlation between *R*_g_ and *c*_sat_ (***Tesei et al., 2021***; ***Bremer et al., 2022***), but the case of V3 and V4 show that certain sequence features can break this symmetry even with- out changing the amino acid composition, and that this is captured by CALVADOS. Examining the sequence features of V3 and V4, we note that V4 has a greater value of *κ* (indicating that negatively and positively charged residues are not well mixed) (Fig. 4a), while the high value of *cv*_aro_ in V3 show that the aromatic residues are highly segregated (Fig. 4b); a feature that has previously been corre- lated with an increased propensity to form amorphous aggregates (***Martin et al., 2020***). Whether these or other sequence features cause the ‘symmetry breaking’ between *R*_g_ and *c*_sat_ for V3 and V4 will be an interesting topic for future analyses.

### Designed variants in the context of the human disordered proteome

The results described above show that we can design IDPs with specific levels of compaction and that charge segregation emerges as an important determinant of compaction of the designed sequences. This result is in line with previous observations from theory, simulation and experiments (***Das and Pappu, 2013***; ***Sherry et al., 2017***; ***Choi et al., 2020***). Recently, we have performed simulations of all IDPs from the human proteome (the IDRome), and found that chain compaction of this broad range of natural sequences is governed by a complex interplay between average hydrophobicity, net charge and charge patterning (***Tesei et al., 2023***). Motivated by these observations we examined the results of the sequences generated by our design algorithm in the context of the properties of natural disordered sequences in the human proteome.

The first aspect which we examined was inspired by our observation that we could generate more compact variants of *α*Syn, A1-LCD and LAF-1-RGG, but not expand these proteins much (Fig. 2). As discussed above, we speculated that this observation was due to the fact that the charged residues in these proteins are already well-mixed so that it is easier to compact them by segregating positive and negative charges than to expand them by further mixing these charged residues. Similarly, we hypothesized that the small number of charged residues in FUS-PLD was the cause of the inability to change the compaction substantially. These observations led us to hypothesize that it would be possible to increase the compaction of natural proteins with stronger charge segregation. We therefore turned to calculations of the *z*(*o*_+−_) score, which is analogous to the *κ* score for charge segregation, but is defined in a way that makes it more appropriate for comparisons across sequences of different lengths and compositions (***Cohan et al., 2021***). We thus examined the distribution of *z*(*o*_+−_) scores across the human IDRome (***Tesei et al., 2023***) and find that, for example, A1-LCD has a well-mixed arrangement of charges as indicated by *z*(*o*_+−_) ::: 0 (Fig. 6a).

**Figure 6.**
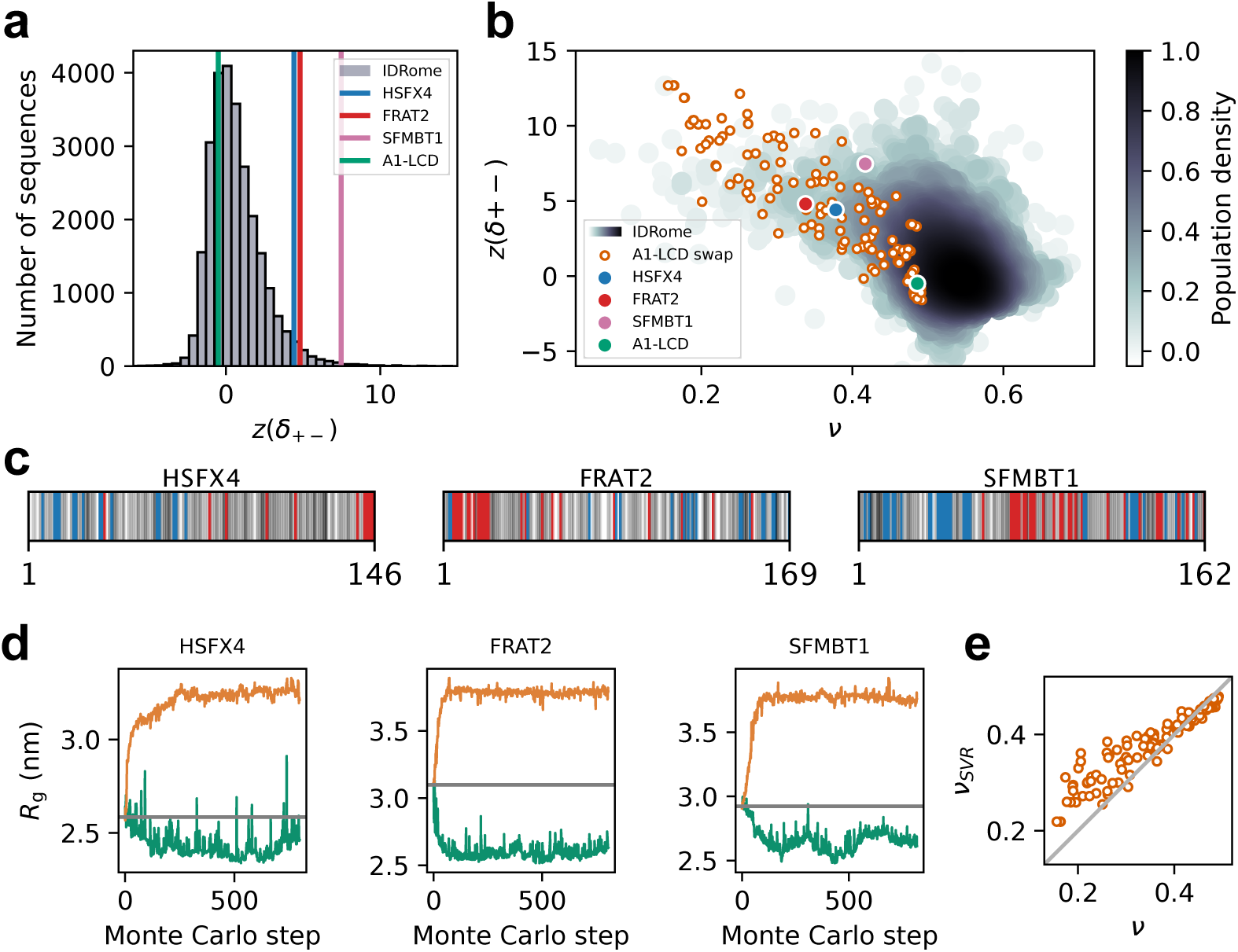
Designed swap variants in the context of the IDRome. (a) Histogram of the sequences in the IDRome grouped based on their charge clustering. We use *z*(*o*_+−_) to compare the degree of charge clustering for sequences of different lengths and composition, with high values of *z*(*o*_+−_) indicating high segregation (***Cohan et al., 2021***). *z*(*o*_+−_) for the wild-type A1, HSFX4, FRAT2, SFMBT1 are indicated respectively in green, blue, red and pink. (b) Comparison of 120 swap variants of A1-LCD (orange) with the IDRome by compaction (*v*) and charge clustering (*z*(*o*_+−_)). (c) Diagram of the sequences of disordered regions in HSFX4, FRAT2 and SFMBT1 that we extracted from the IDRome as representative naturally occurring IDPs that show strong charge clustering. Negative and positive charges are coloured respectively in red and blue. The neutral residues are coloured by a grey scale that reflects their hydrophobicity (corresponding to the *-1* parameter in CALVADOS), with the least hydrophobic residues in white and the most hydrophobic residues in black. (d) Design of more expanded and more compact swap variants starting from the wild-type sequences of the disordered domains of HSFX4, FRAT2 and SFMBT1. (e) Comparison of *v* calculated from MD simulations (with CALVADOS 2 (***Tesei and Lindorff-Larsen, 2022***)) and predicted via an SVR machine-learning model (*v*_SVR_) (***Tesei et al., 2023***) for 120 representative A1-LCD variants.

To examine whether charge patterning and compaction of the designed variants reflect the same rules as for natural proteins we turned to the calculation of scaling exponents (*v*) as a length-independent measure of compaction. For a so-called ‘ideal-chain’ polymer, protein–protein, pro- tein–water, and water-water interactions are balanced, and *v* = 0.5; smaller values of *v* indicate more compact sequences, and an expanded, excluded-volume random-coil has *v* ::: 0.6. We calcu- lated *v* for the designed A1-LCD variants and find that they follow the overall general relationship between charge segregation (*z*(*o*_+−_)) and sequence compaction (*v*) observed for natural proteins (Fig. 6b).

To explore these aspects further, we selected three naturally occurring human IDPs (the disor- dered domains of HSFX4, FRAT2 and SFMBT1) whose compaction can be explained by their strong segregation of positively and negatively charged residues (Fig. 6c). Building on our hypothesis of why we could not expand the well-mixed sequences of *α*Syn, A1-LCD and LAF-1-RGG (Fig. 2), we asked whether we could design sequences resulting in more expanded conformational ensembles if we started from these charge segregated sequences. Indeed, when we applied our design algorithm with the wild-type sequences of HSFX4, FRAT2 and SFMBT1 as starting points, we were able to obtain substantially more expanded sequences as well as also modestly more compact sequences (Fig. 6d). Together, these results support the notion that—for fixed sequence composition—modulation of the distribution of the positively and negatively charged residues is a key determinant of compaction and our ability to change this.

While charge segregation is important for fixed sequence composition, we previously found a more complex interplay between a wider range of sequence properties and chain compaction (***Tesei et al., 2023***). These observations in turn enabled us to train a support vector regression (SVR) machine-learning model to predict scaling exponents from sequences (*v*_SVR_). Given that the SVR model was trained on natural sequences, we asked how well our machine learning model was able to predict chain compaction for designs that have properties that are less common in natural sequences. Overall, we find a high correlation between predicted (*v*_SVR_) scaling exponents and those obtained directly from simulations (*v*) of the 120 A1-LCD variants (Fig. 6e). The aver- age absolute error of the predictions (14%) is somewhat greater than the value found across the IDRome (2.3%; ***Tesei et al. (2023)***), though these values are not fully comparable due to the differ- ent ranges of scaling exponents in the two data sets. We note that defining and calculating scaling exponents is most robust for proteins that behave more like long homopolymers, and that the specific structural properties in the most compact sequences make the average scaling exponent less representative of the conformational ensemble.

## Conclusions

Intrinsically disordered proteins and regions play important roles in a range of biological processes and convey functions that complement those of folded proteins. Thus, the ability to design disor- dered sequences could substantially expand our ability to design proteins with novel functions and properties, in the same way as biology exploits combinations of order and disorder. Combinations of experiments and simulations has led to an improved understanding of the conformational prop- erties of IDPs, which in turn has enabled improved models to generate conformational ensembles directly from sequence via molecular simulations (***Vitalis and Pappu, 2009***; ***Shea et al., 2021***). These models have enabled previous work on design of IDPs (***Zeng et al., 2021***; ***Lichtinger et al., 2021***) and genome-wide studies of sequence-ensemble relationships (***Tesei et al., 2023***; ***Lotthammer et al., 2023***).

Here, we describe a general approach for designing IDPs that exploits a computationally ef- ficient simulation model. Our design algorithm is based on MCMC sampling of sequence space, where each sequence is structurally characterized by combining CALVADOS-based MD simulations (***Tesei et al., 2021***) and alchemical free-energy calculations (***Shirts and Chodera, 2008***). The MCMC sampling guides the sequence towards a design target, and uses the MD simulations and alchem- ical calculations to predict the conformational ensembles of candidate sequences. Together, this leads to an efficient algorithm that we have successfully used to generate a wide range of se- quences with diverse structural features.

We selected five variants of A1-LCD for experimental characterization and find good agreement between experiments and simulations both in terms of the target property (compaction) as well as the propensity of the sequences to undergo phase separation. These findings are in our view important. First, we selected A1-LCD because it is one of the more compact IDPs that have been characterized experimentally, and thus making it even more compact is non-trivial. Second, we restricted our optimization algorithm to maintain sequence composition, and show that we can find substantially more compact sequences even with this restriction. Third, the high correlation between the experimental and calculated radii of gyration demonstrates that CALVADOS remains accurate even for highly unnatural sequences whose properties are well outside those it has pre- viously been trained and benchmarked on. This is a strong validation of our approach of using a physics-based model to drive the sequence design algorithm. We note, however, that the CALVA- DOS force field we used could have been readily reparameterized to improve predictions of single- chain compaction, in case our experiments had revealed discrepancies with simulation predictions (***Norgaard et al., 2008***; ***Tesei et al., 2021***). Fourth, we show that our designs not only match the experiments for the design target (compaction), but also have phase separation properties that generally match the predictions from simulations. We note, however, that V5 appears to be an outlier since its experimental *c*_sat_ value is lower than the prediction from CALVADOS and deviates from the observed trend of increasing *c*_sat_ with increasing *R*_g_. The origin of the discrepancy for the *c*_sat_ value is unclear and we note again that we accurately predict the *R*_g_ of V5.

In addition to developing an algorithm to design IDPs with different levels of compaction, our work also sheds light on sequence-ensemble relationships that can help us understand how natu- ral evolution shapes IDPs. We found that we could generate more compact structures for proteins with the same composition as *α*Syn, A1-LCD and LAF-1-RGG, but not for FUS-PLD, and that we could not generate substantially more expanded conformations based on any of these compositions. Our results show that these effects are mainly due to the number and patterning of charged residues in these proteins. Thus, while global sequence composition may be an important factor in the evolution of IDPs (***Hansen et al., 2006***; ***Tompa and Fuxreiter, 2008***; ***Moesa et al., 2012***) our results support the notion that patterning also plays a key role. The results from these analysis are in line with previous bioinformatics analyses that show that most natural IDPs have relatively high mixing of positively and negatively charged residues (***Holehouse et al., 2017***). Nevertheless, we and others have previously shown that some natural IDPs are compact due to strong segregation of positively and negatively charged residues (***Das and Pappu, 2013***; ***Sawle and Ghosh, 2015***; ***Tesei et al., 2023***; ***Lotthammer et al., 2023***), and we show that for sequences such as the disordered domains of HSFX4, FRAT2 and SFMBT1 we can indeed generate more expanded sequences by disrupting this charge patterning. Whether the high mixing of charged residues is due to entropic effects of many tolerated mutations in IDPs (***Nilsson et al., 2011***; ***Schlessinger et al., 2011***; ***Pajkos et al., 2012***; ***Forman-Kay and Mittag, 2013***) or is due to effects e.g. on solubility or preventing erroneous interactions is an interesting question for future studies.

Looking ahead, our results show that the accuracy of CALVADOS appears to extrapolate also outside the realm of the natural proteins, and variants thereof, on which the model was trained. This suggests that even more extensive sampling of sequence space might be useful. While our MCMC-based approach enables a fine-grained and substantial sampling of the sequence space, it may be combined with or replaced by other approaches to guide the sequence design. We and others have recently shown that it is possible to encode the sequence-ensemble relationships from coarse-grained simulations in machine learning methods (***Tesei et al., 2023***; ***Lotthammer et al., 2023***; ***Chao et al., 2023***); we suggest that such methods for predicting properties from sequences may be used together with, for example, reinforcement learning (***Angermueller et al., 2020***; ***Wang et al., 2023***) or Bayesian optimization (***Yang et al., 2022***) to explore sequence space even more efficiently. This would in particular be important when designing for structural observables that are more complex than single-chain compaction, where simulations could be more expensive and alchemical free-energy calculations might be less efficient. Indeed, our algorithm is general and can be applied to design for other structural features than compaction, and can be adapted to other ways of sampling sequence space. The range of applications can therefore be extended to studies focused on understanding the effect of the patterning of specific residues or groups of residues, or to designing for e.g. binders for disordered therapeutic targets.

In summary, we have developed, applied and validated an algorithm for designing disordered sequences with specified conformational properties. We show that we can design IDPs with substantially increased compaction even with fixed amino acid composition, and find that our algorithms mostly exploits the relationship between charge patterning and compaction. We also explain why some sequences are difficult to expand when the positively and negatively charged residues are well-mixed. Our experimental validation highlights the accuracy of the coarse-grained model with prospective testing of novel sequences. Together, our results show that it is now possible to design sequences of disordered proteins, thus expanding our toolbox for designing proteins with novel or improved functions.

## Methods

### Markov chain Monte Carlo sampling for IDP design

We employed a MCMC algorithm to generate sequences of IDPs. We here targeted the compaction of the chain (as quantified by the *R*_g_) and kept the composition constant during the sequence sampling by using swaps of a randomly selected pair of residues as our MCMC move. We evaluated the *R*_g_ of the new sequence, either by running an MD simulation or by reweighting (see below), and used the Metropolis-Hastings criterion to evaluate the probability of acceptance (*A_k_*_−1---_*_k_*):

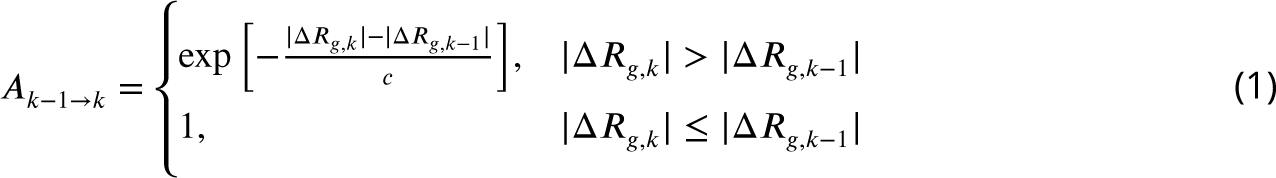

Here, li*R*_g,k_ is the cost function that quantifies the absolute difference between the *R*_g_ of the sequence at the MCMC step *k* and a target *R*_g_ (li*R*_g,k_ = *R*_g,k_ −*R*_g,target_), and *c* is a control parameter. *R*_g,target_ is set to 0 nm to design for more compact IDPs and to 10 nm to design for more expanded IDPs. The starting value for *c* is 0.014, corresponding to *A_k_*_−1---_*_k_*=0.5 for li*R_g,k_* − li*R_g,k_*_−1_ =0.01 nm. We apply simulated annealing using an approach where *c* is decreased by 1% every 2*/* MCMC steps, where */* is the number of amino acids in the IDP sequence.

Although in this work we focus on the specific application of generating variants with fixed amino acid composition, the algorithm and our software accommodates other user-specified MCMC moves (e.g. single- or multi-site amino acid substitutions, substitutions restricted to specific positions and specific residue types). Furthermore, other observables that can be calculated from the simulations can be used as design target. A scheme of the design algorithm is shown in Fig. S10.

### Molecular dynamics simulations

We ran coarse-grained molecular dynamics simulations using the CALVADOS M1 (***Tesei et al., 2021***) *C_a_* -based model. Instead, when comparing *v* from simulations to *v* predicted with the SVR model, we used the CALVADOS 2 (***Tesei and Lindorff-Larsen, 2022***) model since the SVR model was trained on CALVADOS 2 simulations. Single chain simulations in the design algorithm were run for 500 ns with a 10 fs time step. Simulation conditions were set to reproduce 298 K, 150 mM ionic strength and pH 7. Other single chain simulations that are not in the context of the design were run for 1 *µ*s and, when simulations are compared to experiments, at the experimental conditions.

Multi-chain simulations to study the PS propensity of the A1-LCD variants were performed in slab geometry with the CALVADOS M1 model. One hundred chains were assembled in a simulation box 150 nm long and with a cross-section of 15 nm×15 nm. Multi-chain simulations were run for 20 *µ*s. For multi-chain simulations of experimental constructs, three replicates were run for a total simulation time of 120 *µ*s (one replicate 20 *µ*s long and two replicates 50 *µ*s long).

The cut-off used for nonbonded non-ionic interactions was 4 nm for single-chain simulations and 2 nm for multi-chain simulations (***Tesei and Lindorff-Larsen, 2022***). Charge-charge interactions were truncated and shifted at a cut-off of 4 nm in all simulations.

### Alchemical free-energy calculations with MBAR

When proposing a new sequence, the design algorithm attempts to predict the *R*_g_ by reweighting simulations generated at previous steps of the MCMC algorithm using the Multistate Bennett Acceptance Ratio (MBAR) method (***Shirts and Chodera, 2008***). Since the simulations are performed with a *C_a_* -based coarse-grained model, changing the amino acid type in a position of the sequence simply means changing the force field parameters and possibly the charge of the bead representing the residue at that position. Thus, it is easy to evaluate the per-frame potential energy of a new sequence of conformations sampled with another protein sequence. MBAR takes as input an energy matrix defined by frames coming from *n* simulations of different sequences (MBAR pool) and the potential energy functions from each sequence. We calculate the potential energies of the frames of the simulations for a new sequence proposed by the MCMC algorithm, and use MBAR to obtain the Boltzmann weights to estimate the weighted average of the *R*_g_ of the new sequence without running a new simulation.

The reweighting is most accurate when there is substantial overlap between the potential energy functions of the simulations in the MBAR pool and that of the new sequence. We quantify how much the energies of the frames from the simulations in the MBAR pool are compatible with the potential energy function of the new sequence by calculating the number of effective frames (N_eff_) that contributes to the averaging:

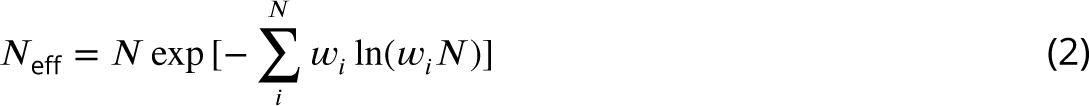

where *N* is the total number of frames from the simulations in the MBAR pool and *w_i_* is the weight of the *i^th^* frame obtained from MBAR to calculate the *R*_g_ of the new sequence. By generating test data sets where we compare the simulated *R*_g_ with the predicted *R*_g_ from MBAR weights, we assessed the relationship between *N*_eff_ and the accuracy of the predicted *R*_g_ (Fig. S4). In light of this analysis, we set a threshold for *N*_eff_ to 20000. When the weights obtained by MBAR result in a *N*_eff_ below this threshold, the algorithm initiates a new simulation and uses the *R*_g_ from this simulation when evaluating the acceptance probability.

The ability to estimate the *R*_g_ of new sequences by reweighting makes the design algorithm more efficient as it decreases the number of MD simulations that are needed. Due to the large size of the energy matrix, we still need to keep the number of simulations in the MBAR pool relatively low, so that the calculations are efficient. With a test data set, we also assessed how the efficiency of the algorithm would change varying the size of the MBAR pool. In general, the larger the pool, the less simulations are required by the algorithm (*i.e.* it occurs less frequently that the *N*_eff_ drops below 20000). In light of these observations, we set the maximum size of the MBAR pool to 10 (Fig. S4). When the size of the pool is at its maximum and the *N*_eff_ drops below the threshold, a new simulation is performed and added to the pool, while the oldest simulation is discarded from the MBAR pool.

### Small-angle X-ray scattering

SAXS (Fig. S11 and Table S2) was performed at BioCAT (beamline 18ID at the Advanced Photon Source, Chicago) with in-line size exclusion chromatography (SEC-SAXS) to separate sample from aggregates, contaminants and storage buffer components, thus ensuring optimal sample quality (Fig. S12) as previously reported (***Bremer et al., 2022***; ***Martin et al., 2020***, ***2021***). Samples were loaded onto a Superdex 75 Increase 10/300 GL column (Cytiva), which was run at 0.6 mL/min by an AKTA Pure FPLC (GE) and the eluate, after passing through the UV monitor, was flown through the SAXS flow cell. The flow cell consisted of a 1.0 mm ID quartz capillary with ∼20 *µ*m walls. All protein solutions were measured at room temperature in 20 mM HEPES (pH 7.0), 150 mM NaCl, 2 mM DTT. A co-flowing buffer sheath was used to separate the sample from the capillary walls, helping prevent radiation damage (***Kirby et al., 2016***). Scattering intensity was recorded using an Eiger2 XE 9M (Dectris) detector which was placed 3.685 m from the sample giving us access to a *q*-range of 0.0029–0.42 Å^−1^. 0.5 s exposures were acquired every 1 s during elution and data were reduced using BioXTAS RAW 2.1.4 (***Hopkins et al., 2017***). Buffer blanks were created by averaging regions flanking the elution peak and subtracted from exposures selected from the elution peak to create the *J*(*q*) vs *q* curves (scattering profiles) used for subsequent analyses. RAW was used for buffer subtraction, averaging, and Guinier fits. Scattering profiles were additionally fit using an empirically derived molecular form factor (MFF) (***Riback et al., 2017***) (used to calculate the experimental *R*_g_ values in Fig. 5).

### Diffusion Ordered NMR Spectroscopy

We carried out diffusion ordered spectroscopy (DOSY) experiments (***Wu et al., 1995***) at 307 K to measure translational diffusion coefficients for WT A1-LCD and the V1 variant, by fitting intensity decays of individual signals selected between 0.5 ppm and 2.5 ppm (***Leeb and Danielsson, 2020***) with the Stejskal-Tanner equation (***Stejskal and Tanner, 1965***). We used 1,4-dioxane (0.10% v/v) as internal reference for the *R*_h_ (2.27 ± 0.04 Å, (***Tranchant et al., 2023***)). We acquired 80 scans for A1-LCD and 480 scans for V1. Spectra were recorded on a Bruker 600 MHz spectrometer equipped with a cryoprobe and Z-field gradient, and were obtained over gradient strengths from 5 to 95% (32 points) for A1-LCD and from 5% to 75% (16 points) for V1 (*γ* = 26752 rad s^−1^ Gauss^−1^) with a diffusion time (li) of 50 ms and gradient length (*δ*) of 6 ms. Translational diffusion coefficients were fitted in Dynamics Center v2.5.6 (Bruker) and were used to estimate the *R*_h_ for the proteins (***Prestel et al., 2018***), with error propagation using the diffusion coefficients of both the protein and dioxane.

### Data and code availability

Data and code used and produced by this study are available on GitHub. MD simulations of 120 A1-LCD variants and of the six experimental constructs of A1-LCD variants and wild-type, both as single-chain and multi-chains in slab geometry, are available on the Electronic Research Data Archive. SAXS data are deposited in SASDB (***Kikhney et al., 2020***) (Table S2).

### Author contributions

F.P, T.M. and K.L.-L. designed the study. F.P, G.T. and K.L-L. handled all computational and theoretical aspects. F.P. and A.B. expressed and purified proteins, measured *c*_sat_ and acquired DIC microscopy images. C.R.G. measured NMR data. F.P. and C.R.G. analyzed NMR data. J.B.H. measured SAXS data. F.P., A.B. and J.B.H. analyzed SAXS data. F.P. and K.L.-L. analyzed the data and wrote the paper with input from all authors.

## Acknowledgments

We thank Wade Borcherds and Emil Tranchant for helpful discussions, and George Campbell for assistance with DIC microscopy. This work was supported by the Lundbeck Foundation BRAINSTRUC structural biology initiative (R155-2015-2666, to K.L.-L.) and the PRISM (Protein Interactions and Stability in Medicine and Genomics) centre funded by the Novo Nordisk Foundation (NNF18OC0033950, to K.L.-L.). We acknowledge access to computational resources from the Danish National Super- computer for Life Sciences (Computerome). This work was supported by the US National Institutes of Health through grant R01NS121114 (T.M.), the St. Jude Research Collaborative on the Biology and Biophysics of RNP granules (T.M.), and the American Lebanese Syrian Associated Charities (to T.M.). We acknowledge use of the Cell and Tissue Imaging Center - Light Microscopy Facility at St. Jude Children’s Research Hospital. This research used resources of the Advanced Photon Source, a U.S. Department of Energy (DOE) Office of Science User Facility operated for the DOE Office of Science by Argonne National Laboratory under Contract No. DE-AC02-06CH11357. BioCAT was supported by grant P30 GM138395 from the National Institute of General Medical Sciences of the National Institutes of Health. The content is solely the responsibility of the authors and does not necessarily reflect the official views of the National Institute of General Medical Sciences or the National Institutes of Health.

