## Supporting Figures, Text and Tables for "Design of intrinsically disordered protein variants with diverse structural properties"

### Supplementary experimental materials and methods

#### Protein constructs

Sequences of wild-type A1-LCD and variants are based on the low complexity domain (residues 186-320) of the human hnRNPA1 (UniProt: P09651; Isoform A1-A). The coding sequences for the variants were synthesized (Thermo Fisher) including a coding sequence for an N-terminal ENLYFQGS TEV protease cleavage site and 5' and 3' attB sites for Gateway cloning. The sequences were recombined via LR reactions into the pDEST17 vector (Thermo Fisher), which includes an N-terminal 6xHis-tag coding sequence. After expression, we cleaved of the N-terminal 6xHis-tag using TEV protease, leaving only an additional GS sequence at the N-terminus.

#### Protein expression and purification

A1-LCD variants were expressed and purified as previously reported for similar constructs (***Martin et al., 2020***; ***Bremer et al., 2022***). The *E. coli* BL21 (DE3) pLysS strain was used for expression and grown in ZYM5052 auto induction media at 37°C for 24 hours. Cell pellets were recovered by centrifugation and resuspended in 50 mM MES pH 6.0, 500 mM NaCl, 20 mM 2-mercaptoethanol. Cell lysis was achieved via sonication. Cell lysates were centrifuged to collect inclusion bodies, that were resuspended in 6 M GdmHCl, 20 mM Tris pH 7.5, 15 mM imidazole overnight at 4°C. The solutions containing the solubilized inclusion bodies were cleared from cell debris by centrifugation, and supernatants were loaded onto self-packed columns of chelating Sepharose fast flow beads (GE Healthcare) charged with nickel sulfate. The columns were washed with four column volumes of 4 M urea, 20 mM Tris pH 7.5, 15 mM imidazole. Proteins were eluted from the Ni-NTA resin with 4 M urea, 20 mM Tris pH 7.5, 500 mM imidazole. TEV cleavage of the 6xHis-tag was done in 2 M urea, 20 mM Tris pH 7.5, 50 mM NaCl, 0.5 mM EDTA, 1 mM DTT overnight at 4°C. Cleaved protein solutions were loaded onto Ni-NTA columns. The flow-through and wash fractions were collected and concentrated using a 3000 MWCO Amicon centrifugal filter. Finally, samples were transfered in 2 M GdmHCl, 20 mM MES pH 5.5 over a S75 Superdex size exclusion column (GE Healthcare). The molecular weight of the proteins and the purity of samples were confirmed via intact mass spectrometry and SDS- PAGE. Samples were stored in 6 M GdmHCl, 20 mM MES pH 5.5 at 4°C.

**Table S1.** Sequences of the wild type A1-LCD and the five designed variants that we characterized experimentally. The first two residues (GS) are left over by the TEV protease cleavage of the 6xHis-tag.

<table>
<tr><th>Label</th><th>Sequence</th></tr>
<tr><td>WT</td><td>GS|MASASSSQRGRSGSGNFGGGRGGGFGGNDNFGRGGNFSGRGGFGGSRGGGGYGGSGDGYNGFGN<br>DGSNFGGGGSYNDFGNYNNQSSNFGPMKGGNFGGRSSGGSGGGGQYFAKPRNQGGYGGSSSSSSYGS<br>GRRF</td></tr>
<tr><td>V1</td><td>GS|GSGSGGSRGGNKRRRKRRGGSGGYRYSRRGGGFNQGGGFNSSGFFGGMGSGGGSGGGFGNGPSFA<br>GSNNFNGGGGGSAGNFGQYGGRGGPYSGSGGSGSGSNSGQNGGSGNYMGSGYDAFYNSSFNNQSFFG<br>DDD</td></tr>
<tr><td>V2</td><td>GS|GGYGSSQGGFFGGGDAGGNGDGSDFGGGYPSGSNQNSGGFSGYGNDSFQGSAGMFNGFKSASKFS<br>NSGGYGGGGQGNNNGSGGGSSFRNRRRRSNYSGGGSGRGRRYGSNFGGMYGGRSGFGGNGPGRSGFG<br>GSN</td></tr>
<tr><td>V3</td><td>GS|KQGGRGGNRSGSGNGNASGAGGGGRDGGSDGGFDGFDYQFSGGGNPSSQYYGSRGGSGRNSAGG<br>YYFFRNSSGGNGSSGNMNPGNGYFGFSRSGGRGQNRGFFFGGMGGGGFGRSSNFGSYNSSNKSGSGGG<br>GGG</td></tr>
<tr><td>V4</td><td>GS|GSNGGGSQSSGQGYGKSGGNRRRGRGGAGGGFGMGDGSNQYGYGPFRRGSGFNGNGDYANYGG<br>NGDSNNFSNYRGGNSANGNFQSGGGGGFDNGGGSGFGGSFSMSGGSSSGKRRGSGGFFSGRSGSGFGG<br>FYPS</td></tr>
<tr><td>V5</td><td>GS|GFSNMGNGFGGRFGGGRGFSRYSQQFSYYDGGQSSGGNGSSGGFNSYGGYNNGRNGSSFGGAGG<br>GGRSSFGFSGGGGFGADGGYNRFSSGDRNNNGPSKGGGGGNGSGSRGFAGNGSMSDRGNSYGGGPGR<br>QKGS</td></tr>
</table>

### SDS-PAGE

Gel electrophoresis was carried out using NuPAGE 4–12% Bis-Tris gradient gels (Invitrogen). 1x NuPAGE MES SDS Running buffer (Invitrogen) was used to run gels. After the run, gels were washed with water and stained with SimplyBlue SafeStain (Thermo Fisher Scientific) before destaining with water. PageRuler Plus Prestained protein ladder (Thermo Fisher Scientific) was used as a molecular weight reference.

### Buffer exchange

To remove the denaturant buffer used for storage and transfer the protein to 20 mM HEPES (pH 7.0) we used Zeba$^{TM}$ Spin Desalting Columns (Thermo Fisher Scientific) with 7k MWCO and 0.5 mL volume. After removal of storage solution from the column by centrifugation at 1000 $\times g$ for 1 min, columns were washed three times with 300 $\mu$L of 20 mM HEPES (by centrifugation at 1000 $\times g$ for 1 min). Finally, protein sample is applied to the column and recovered in 20 mM HEPES after a centrifugation. Additional washing steps (3–5) were carried out in Amicon Ultra-0.5 Centrifugal Filter Units to remove residual denaturant.

### Determination of saturation concentrations

Phase separation of protein samples was induced by adding NaCl to a final concentration of 150 mM. The dilute and dense phase were separated via centrifugation (***Milkovic and Mittag, 2020***). The $c_{sat}$ was determined by the absorbance of the dilute phase at 280 nm.

### DIC microscopy

Differential interference contrast microscopy (DIC) images were obtained at room temperature using a Nikon Eclipse Ni Widefield microscope with a 20X objective. Samples were at concentrations slightly above their $c_{sat}$ at room temperature. Phase separation was induced by adding NaCl to the protein stock solution to reach a concentration of 150 mM. 2 $\mu$L of the protein solution were positioned in between two glass coverslips held toghether by 3M 300 LSE high-temperature double-sided tape (0.34 mm) with a window for microscopy cut out.

### Supplementary computational methods

The $R_\mathrm{h}$ for protein conformations was calculated using HullRadSAS (***Fleming et al., 2023***; ***Tranchant et al., 2023***). The ensemble-averaged $R_\mathrm{h}$ was calculated as $1/n^{-1}\sum_i^n(1/R_{\mathrm{h},i})$ (***Choy et al., 2002***; ***Ahmed et al., 2020***), from each conformer $i$ of an ensemble. Sequence clustering was performed with a 65% sequence identity threshold using the CD-HIT software (***Li and Godzik, 2006***; ***Fu et al., 2012***). Calculations of $\omega_{aro}$ and $\kappa$ from sequences were performed using the localCIDER python package (https://github.com/Pappulab/localCIDER), while the $z(\delta_{+-})$ scores for the IDRome sequences and the A1-LCD swap variants was calculated using a modified version of the NARDINI software which allowed us to define a custom threshold for the largest fraction of negatively and positively charged residues below which the program sets $z(\delta_{+-})$ to zero (***Cohan et al., 2021***). We set this threshold to 2.5% to obtain a non-zero $z(\delta_{+-})$ score for A1-LCD and sequences in the IDRome with fraction of charged residues similar to A1-LCD. For the NARDINI analysis of IDRome sequences, we generated $10^5$ randomly shuffled sequences, while for the wild type and variants of A1-LCD, we used $5{\times}10^5$ randomly shuffled sequences.

We calculated error bars on averages calculated from MD simulations using block averaging (https://github.com/fpesceKU/BLOCKING). Calculation of SAXS data from conformations was performed with Pepsi-SAXS (v3.0) (***Grudinin et al., 2017***), using fixed parameters for the contrast of the hydration layer and the effective atomic radius (respectively 3.34 e/nm$^3$ and $1.025 \times r_\mathrm{m}$, where $r_\mathrm{m}$ is the average atomic radius of the protein) (***Pesce and Lindorff-Larsen, 2021***). Prior to calculating the $\chi_r^2$, experimental SAXS curves are rebinned to 158 scattering angles and experimental error bars are rescaled using the Bayesian indirect Fourier transform (BIFT) (***Larsen and Pedersen, 2021***). Both rebinning and error correction were carried out with the BayesApp webserver (https://somo.chem.utk.edu/bayesapp/) (***Hansen, 2012***).

### Supplementary figures and tables

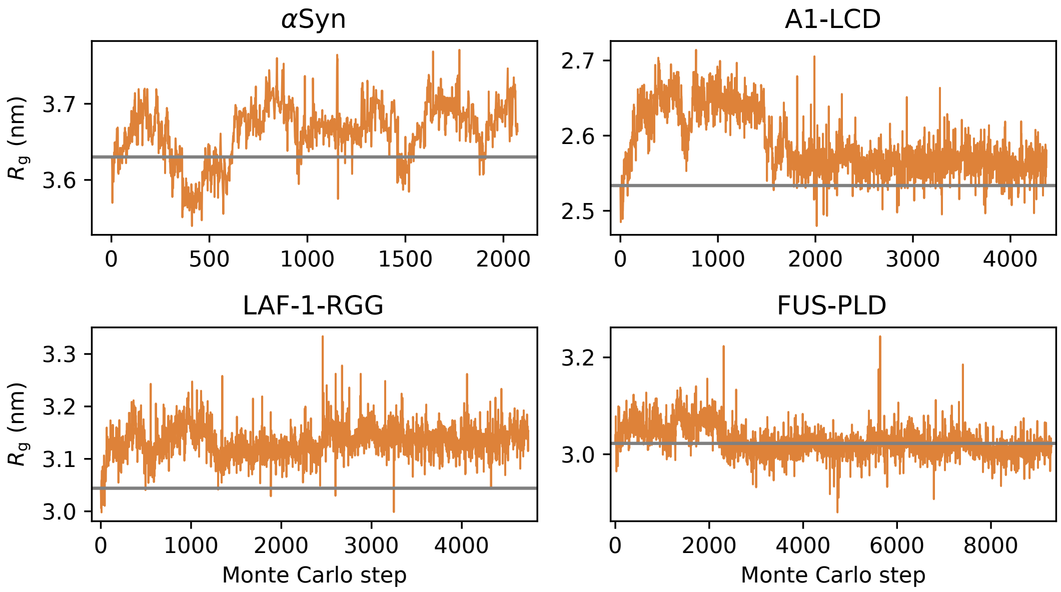


**Figure S1.** Design of more expanded variants for $\alpha$Syn, A1-LCD, LAF-1-RGG and FUS-PLD, starting from the wild-type sequences.

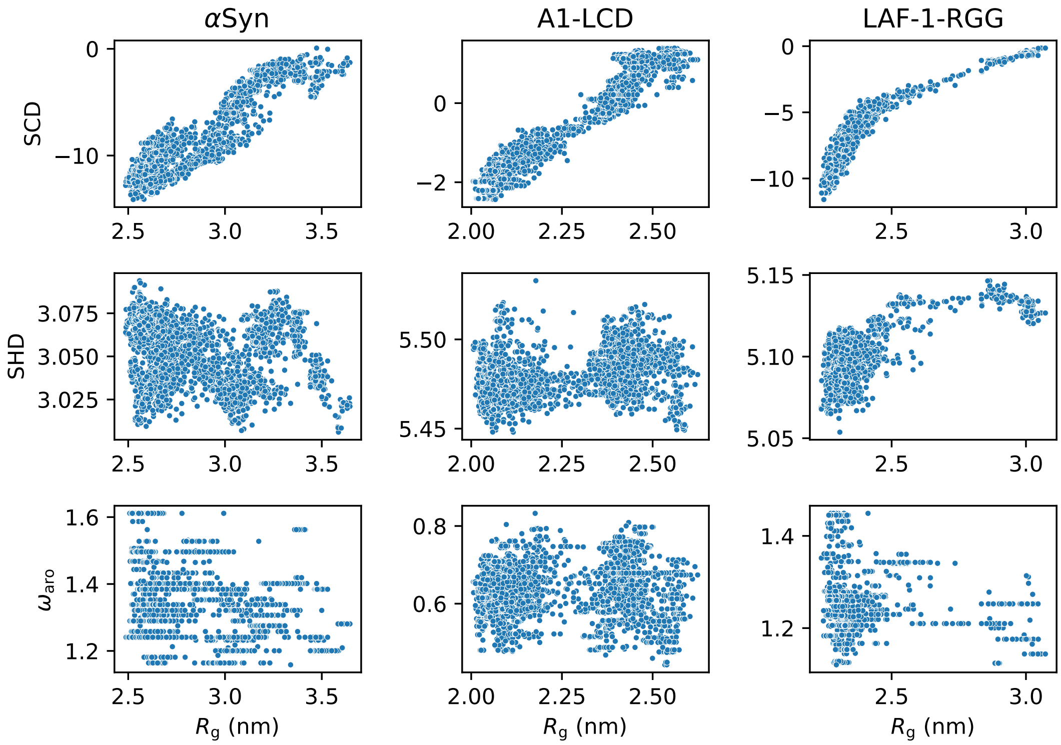


**Figure S2.** Multiple sequence features were calculated from the variant sequences of $\alpha$Syn, A1-LCD and LAF-1-RGG and correlated with the $R_g$. SCD, similarly to $\kappa$, is related to the patterning of charged residues. SHD (sequence hydropathy decoration) quantifies the patterning of hydrophobic residues. $\omega_{aro}$ quantifies the patterning of aromatic residues.

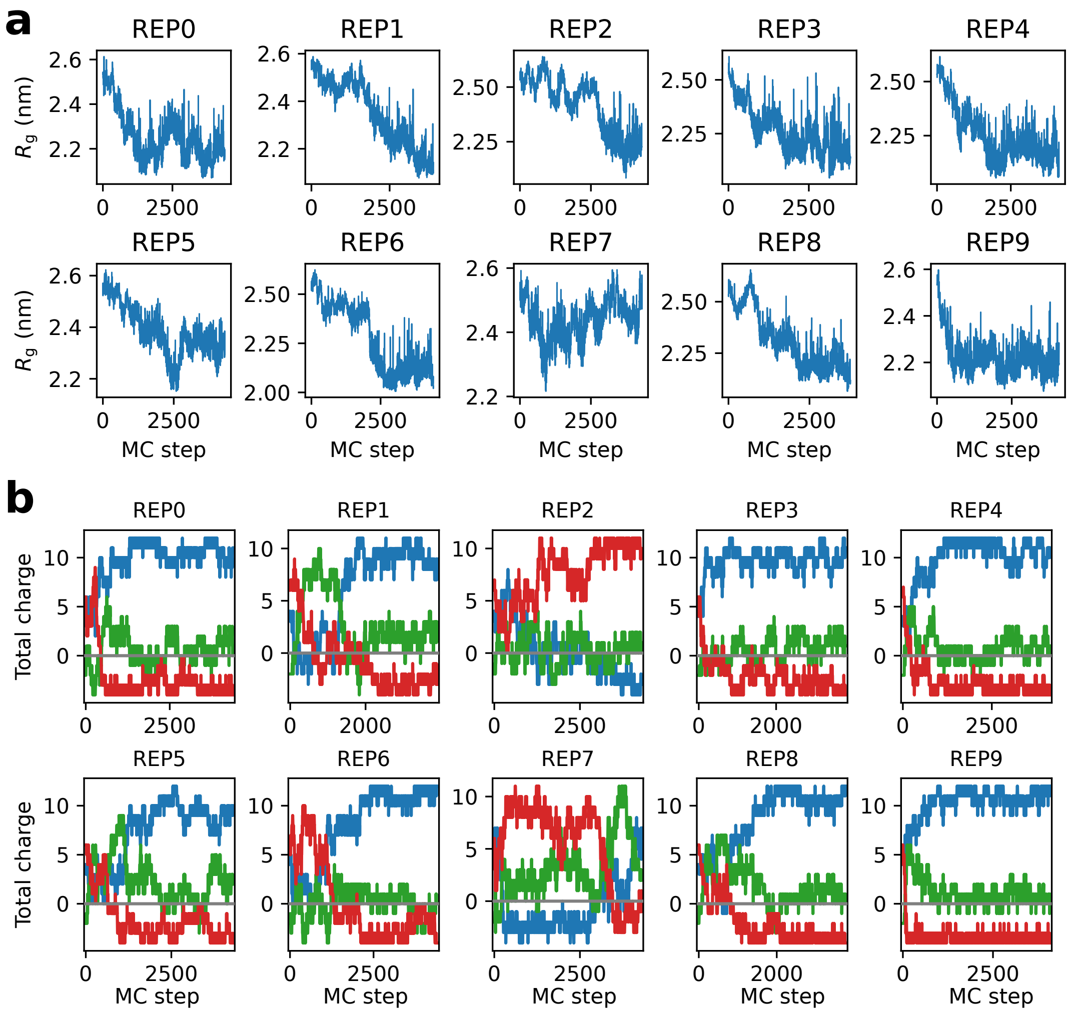


**Figure S3.** We performed ten runs for generating compact variants of A1-LCD. For each replica we show (a) the evolution of the $R_g$ from the generated sequences and (b) the total charge for the N-terminal third (blue), the middle third (green), and the C-terminal third (red) of each sequence.

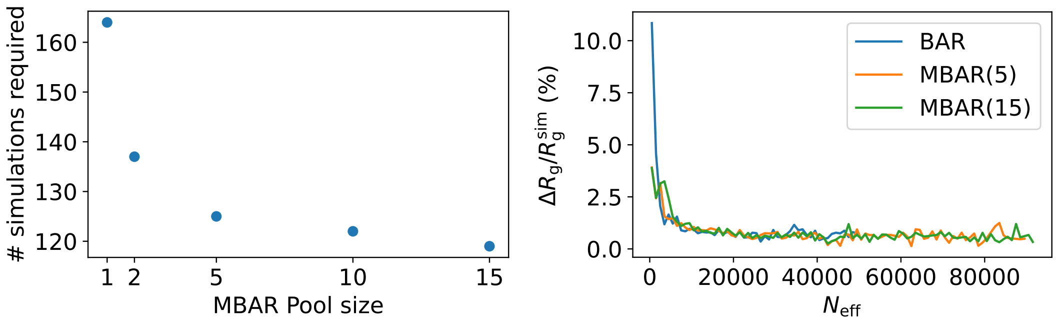


**Figure S4.** To test the accuracy and efficiency of MBAR reweighting, we generated a random sequence of 140 residues and performed 1000 position swaps between two randomly selected residues. We simulated all 1000 sequences and calculate their $R_g$. Then we iterate through the 1000 sequences trying to predict their $R_g$ by reweighting simulations from previous iterations. We vary the maximum size of the MBAR pool and add a new simulation to the pool when the $N_{\mathrm{eff}}$ drops below 10000. Then we compare the reweighted $R_g$ from MBAR with the simulated $R_g$. The left panel shows the number of simulations required by varying the maximum MBAR pool size. The right panel shows the relative absolute difference between reweighted and simulated $R_g$ ($|\Delta R_g|/R_g^{\mathrm{sim}}$) as a function of $N_{\mathrm{eff}}$. For better visualization, we binned the data on the $N_{\mathrm{eff}}$ coordinate (with a bin width of 1000) and plot the average in the bins.

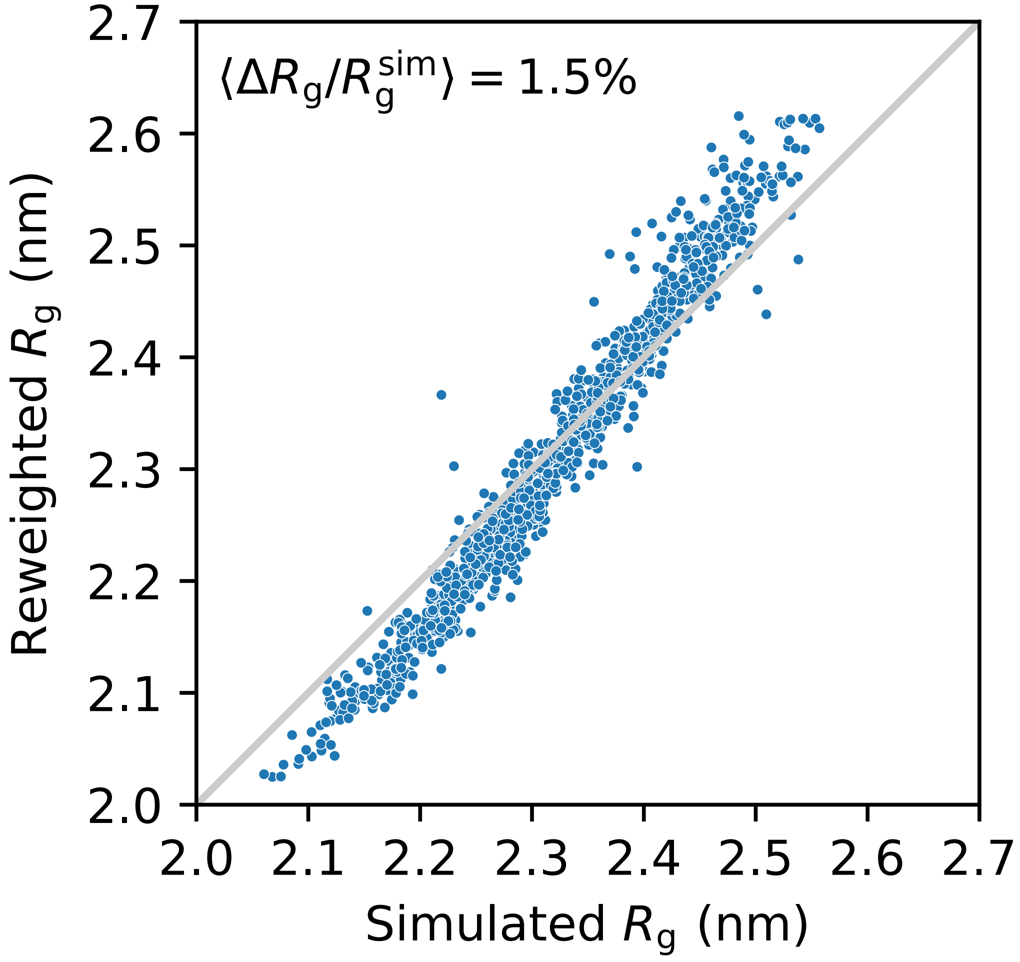


**Figure S5.** For some of the centroids selected from the sequence clustering of the A1-LCD variants the $R_g$ values had been obtained by reweighting. We simulated each of these for 1 $\mu$s to assess the accuracy of the reweighting. The reweighted and simulated $R_g$ values are compared. We observe an average error of 1.5% on the reweighted $R_g$, with a slight bias for the most compact and expanded chains.

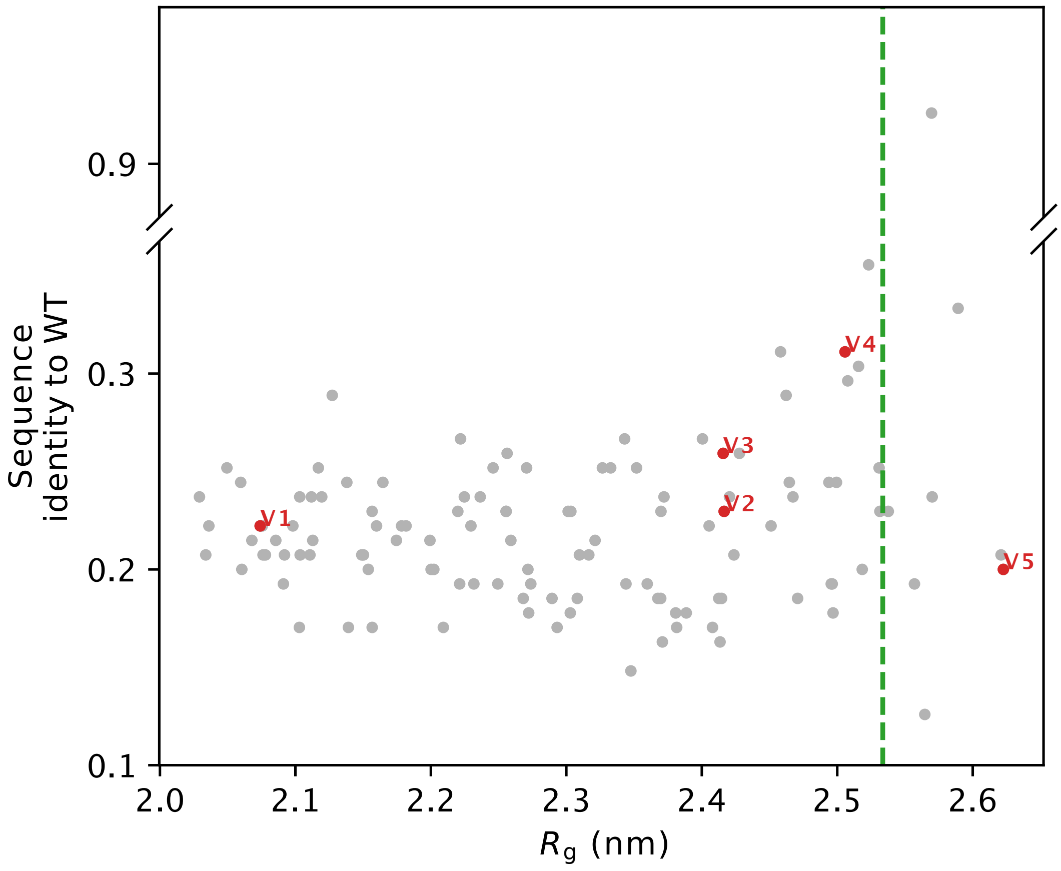


**Figure S6.** Sequence identity to wild-type A1-LCD for the 119 designed A1-LCD variants. Green vertical line correspond to the $R_g$ of wild-type A1-LCD.

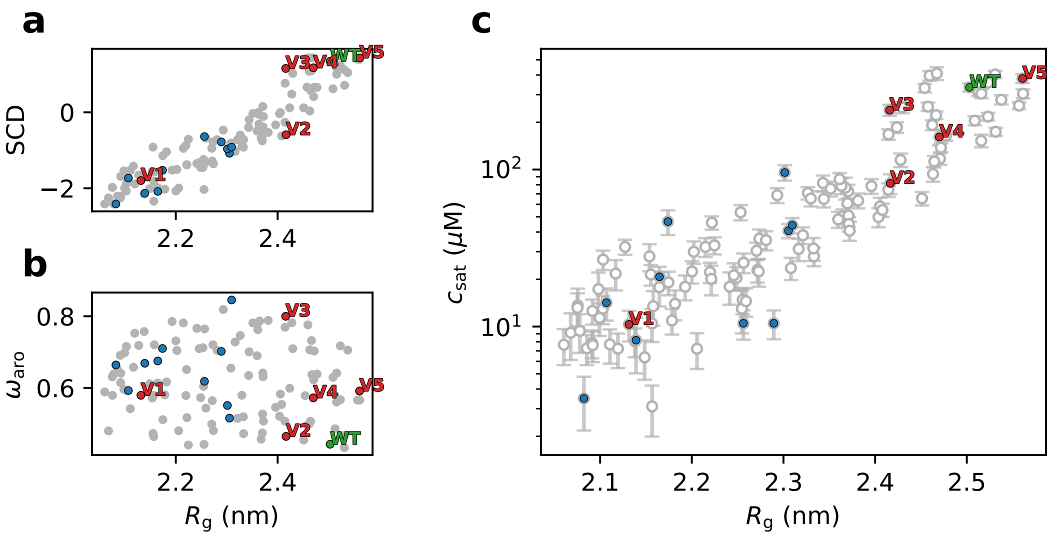


**Figure S7.** Characterization of the 120 variants of A1-LCD. We show the relationship between $R_g$ and (a) SCD, (b) $\omega_{aro}$ (patterning of aromatic residues) and (c) the $c_{sat}$ calculated from simulations of 100 chains in slab geometry. We highlight the wild-type sequence of A1-LCD in green, the five variants that we characterized experimentally in red, and ten variants that did not express in *E. coli* in blue.

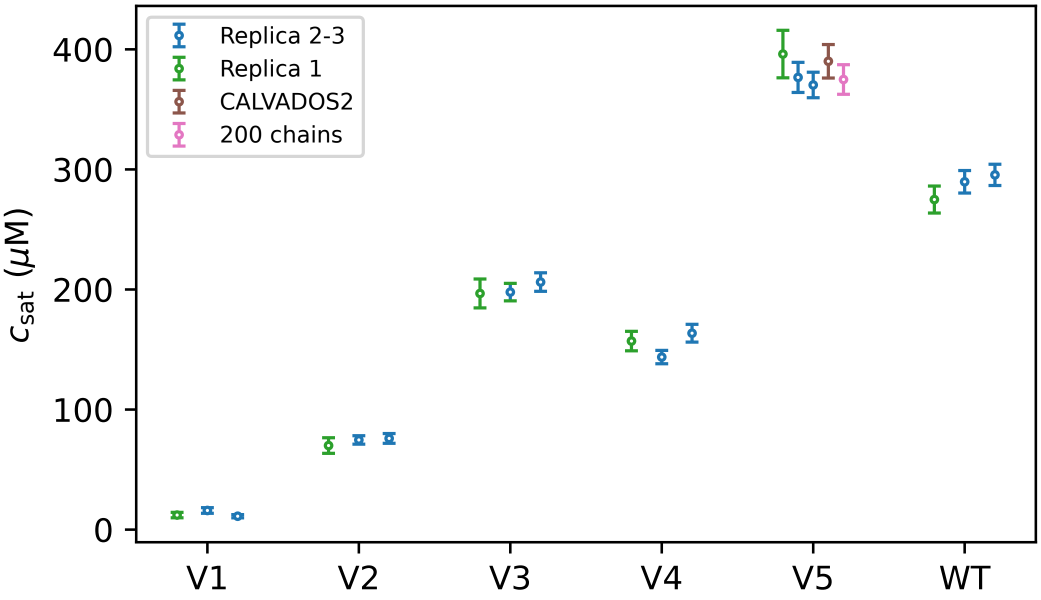


**Figure S8.** $c_{sat}$ values calculated by slab simulations of experimental constructs. Replicas 1,2 and 3 were performed with CALVADOS M1 (***Tesei et al., 2021***) and 100 chains in the simulation box. Replica 1 (green) is 20-$\mu$s long, while replicas 2 and 3 (blues) are 50-$\mu$s long. For V5, we also performed a 20-$\mu$s long simulation with CALVADOS M1 but using 200 chains (pink), and a 20-$\mu$s long simulation with 100 chains but the CALVADOS 2 parameters (brown) (***Tesei and Lindorff-Larsen, 2022***).

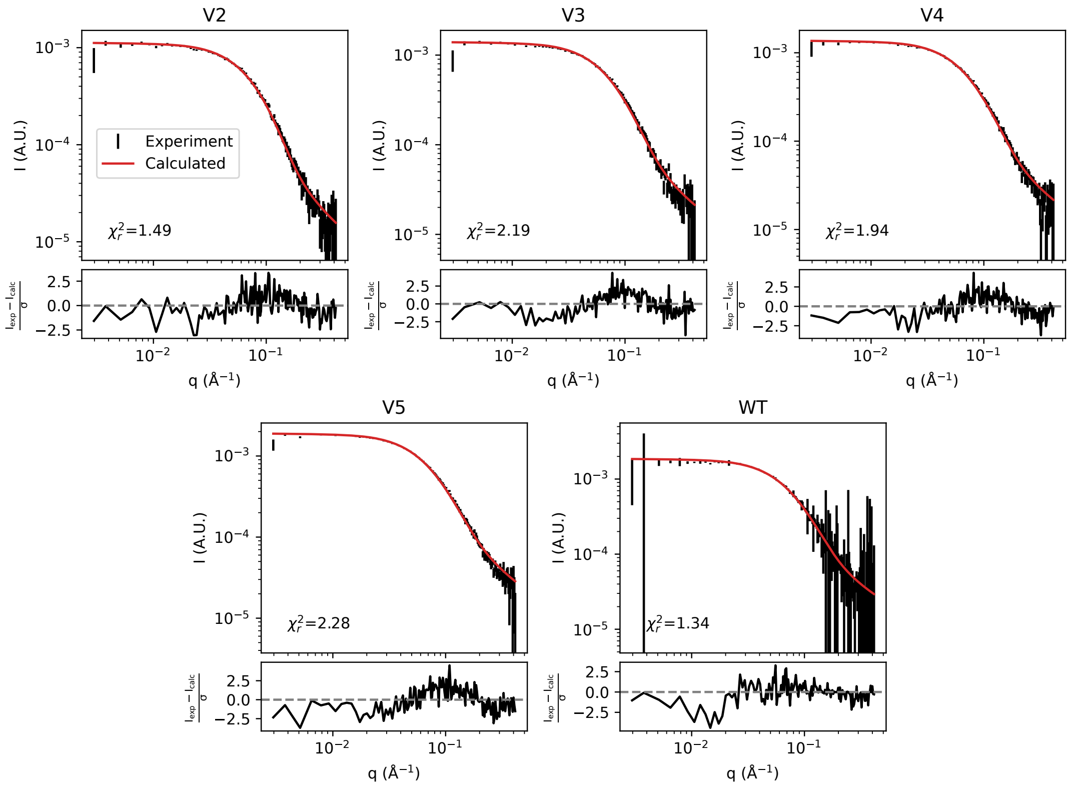


**Figure S9.** Rebinned experimental SAXS data with corrected error bars (black) compared to SAXS curves calculated from simulations.

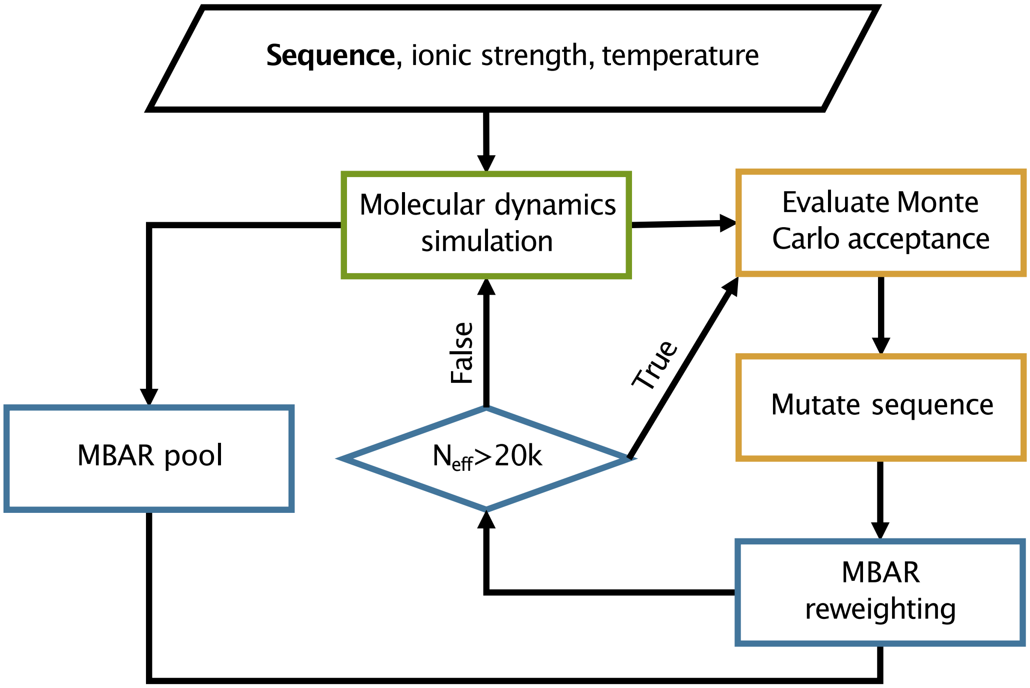


**Figure S10.** Schematic outline of the design algorithm.

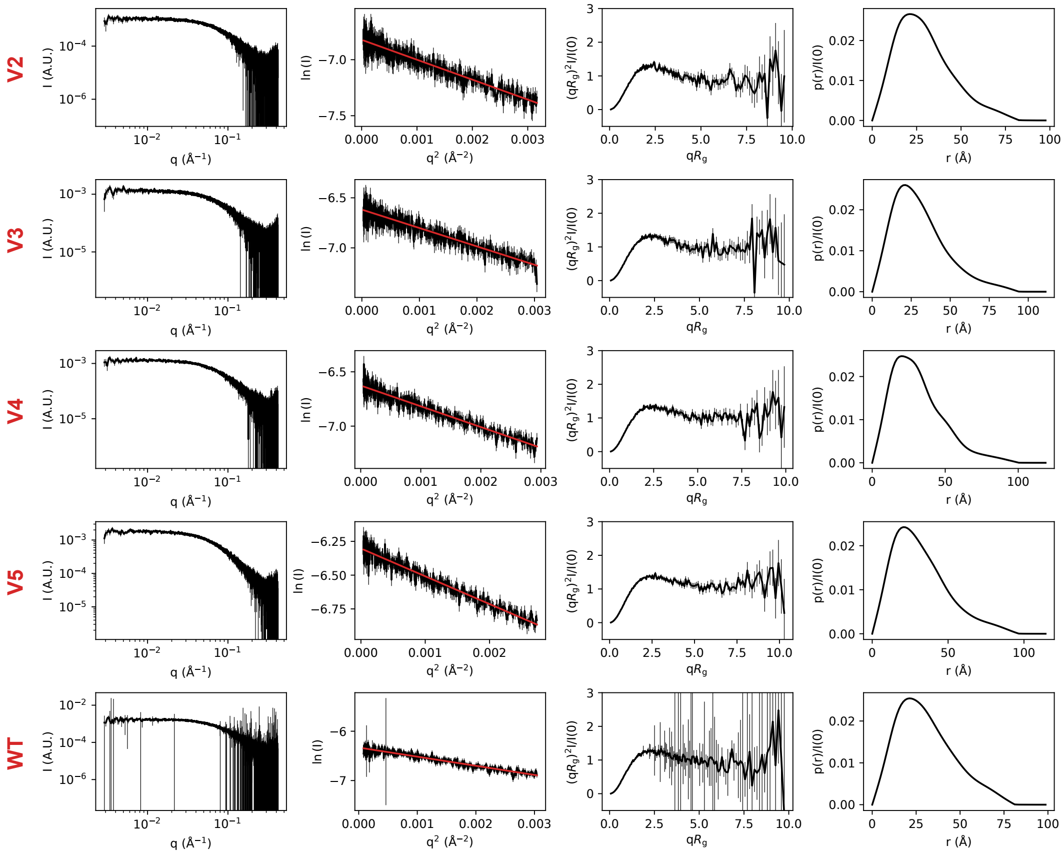


**Figure S11.** SAXS data collected on samples of (from top to bottom rows) V2, V3, V4, V5 and wild-type A1-LCD. From the left to the right column, SAXS profiles are shown with logarithmic scales, as a Guinier plot in the range used for the linear fit (in red) to derive the the Guinier $R_g$, the dimensionless Kratky plot with rebinned SAXS data, and the normalized pair distance distribution function (calculated using BIFT (***Larsen and Pedersen, 2021***)).

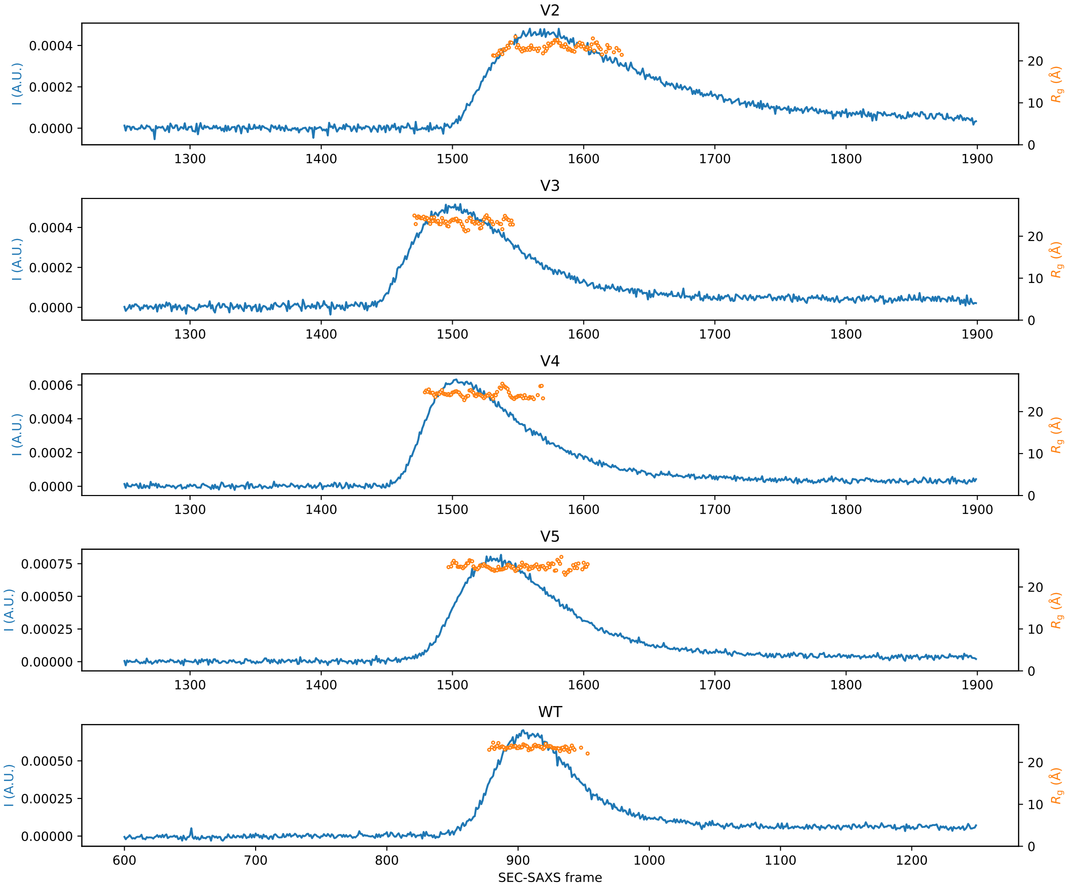


**Figure S12.** In line SEC-SAXS data collection. In blue, we show the mean solvent-subtracted intensity for each SAXS frame collected during sample elution from the SEC column. In orange, we show the Guinier $R_g$ calculated for each SAXS frame.

**Table S2.** SAXS sample, data-collection and analysis for the wild-type A1-LCD and its variants*.

| **(a) Sample details** | | | | | |
|---|---|---|---|---|---|
| | **V2** | **V3** | **V4** | **V5** | **WT** |
| Organism | Artificial | Artificial | Artificial | Artificial | Human |
| Source | | | *E. coli* BL21 (DE3) pLys recombinant expression | | |
| *Sample environment/configuration* | | | | | |
| Solvent composition | | | 20 mM HEPES pH 7.0, 150 mM NaCl, 2 mM DTT | | |
| Sample temperature (K) | | | 298 | | |
| In-beam sample cell | | | 1 mm quartz capillary flow cell | | |
| *Size exclusion chromatography* | | | | | |
| Sample injection concentration (mg/mL) | 2.6 | 2.6 | 2.6 | 2.6 | 2.6 |
| Sample injection volume (mL) | | | 250 | | |
| SEC column type | | | Superdex 75 Increase 10/300 GL column (Cytiva) | | |
| SEC flowrate (mL/min) | | | 0.6 | | |
| **(b) SAXS data collection** | | | | | |
| Data-acquisition/reduction software | | | BioXTAS RAW 2.1.4 | | |
| Source/instrument description | | | BioCAT (Sector 18, APS) | | |
| Measured q-range ($q_{\mathrm{min}}$ – $q_{\mathrm{max}}$) (Å$^{-1}$) | | | 2.90e-03 – 4.17e-01 | | |
| Method for scaling intensities | | | Absolute scaling with glassy carbon | | |
| Exposure time (s) | | | 0.5 | | |
| **(c) SAS-derived structural parameters** | | | | | |
| *Guinier analysis* | | | | | |
| Method(s)/software | | | autorg (ATSAS 3.1.3) | | |
| I(0) | 0.0011 ± 5.4e-06 | 0.0013 ± 6.8e-06 | 0.0013 ± 5.3e-06 | 0.0018 ± 6.6e-06 | 0.0018 ± 8.7e-06 |
| $R_g$ (Å) | 23.1 ± 0.2 | 23.48 ± 0.21 | 23.95 ± 0.17 | 24.84 ± 0.16 | 23.55 ± 0.21 |
| $qR_g$ range | 0.13 – 1.3 | 0.12 – 1.3 | 0.16 – 1.3 | 0.15 – 1.3 | 0.21 – 1.3 |
| Linear fit assessment (*autorg* fidelity) | 1 | 1 | 1 | 0.98 | 0.01 |
| *Pair distance distribution function analysis* | | | | | |
| Method(s)/software | | | BIFT (BayesApp 1.1) | | |
| I(0) | 1.09e-03 | 1.35e-03 | 1.34e-03 | 1.85e-03 | 1.79e-03 |
| $R_g$ (Å) | 23.65 | 24.89 | 25.47 | 26.21 | 24.5 |
| $D_{\mathrm{max}}$ (Å) | 82.56 | 93 | 98.89 | 95.35 | 80.32 |
| P(r) reciprocal-space fit: $\chi_r^2$, p-value | 0.80, 4.40e-04 | 0.77, 4.1e-05 | 0.87, 2.6e-02 | 0.84, 4.5e-03 | 0.73, 4.1e-07 |
| **(d) Scattering particle size** | | | | | |
| Porod volume (Å$^3$) | 16726 | 14254 | 14874 | 22680 | 15360 |
| Theoretical MW (kDA) | | | 13.1 | | |
| SAXS MW (DatBayes)** (kDA), probability | 15.475, 0.45 | 14.825, 0.48 | 14.825, 0.43 | 14.825, 0.39 | 15.475, 0.50 |
| **(e) Modelling (SAXS calculation from molecular simulations)** | | | | | |
| Software | | | Pepsi-SAXS (3.0) | | |
| q-range for calculation (Å$^{-1}$) | | | 2.90e-03 – 4.17e-01 | | |
| Number of frames used | | | 10000 | | |
| Scale factor and offset | | | Fixed to constant in Pepsi-SAXS, then globally fit to experiment by least square | | |
| $\delta\rho$ (e/nm$^3$) | | | 3.34 | | |
| Average atomic radius ($r_m$; Å) | | | 1.58 | | |
| $r_0/r_m$ | | | 1.025 | | |
| $\chi_r^2$ | 1.49 | 2.19 | 1.94 | 2.28 | 1.34 |
| **(f) Data deposition** | | | | | |
| SASDB ID | SASDTK2 | SASDTL2 | SASDTM2 | SASDTN2 | SASDTJ2 |

* Table in accordance with guidelines from ***Trewhella et al.*** (***2017***) and ***Trewhella et al.*** (***2023***)

** (***Hajizadeh et al., 2018***)
